# Culling less fit neurons protects against amyloid-β-induced brain damage and cognitive and motor decline

**DOI:** 10.1101/468868

**Authors:** Dina S. Coelho, Silvia Schwartz, Marisa M. Merino, Barbara Hauert, Barbara Topfel, Colin Tieche, Christa Rhiner, Eduardo Moreno

## Abstract

Alzheimer’s disease (AD) is the most common form of dementia, impairing cognitive and motor functions. One of the pathological hallmarks of AD is neuronal loss, which is not reflected in mouse models of AD. Therefore, the role of neuronal death is still uncertain. Here, we used a *Drosophila* AD model expressing a secreted form of human amyloid-β42 peptide and show that it recapitulates key aspects of AD pathology, including neuronal death and impaired long-term memory. We found that neuronal apoptosis is mediated by cell fitness-driven neuronal culling, which selectively eliminates impaired neurons from brain circuits. We show that removal of less fit neurons delays amyloid-β42-induced brain damage and protects against cognitive and motor decline, suggesting that - contrary to common knowledge - neuronal death may have a beneficial effect in AD.

## Introduction

Multicellular organisms have evolved mechanisms to maintain tissue homeostasis and integrity throughout development and ageing. Besides cell-intrinsic surveillance mechanisms, relative fitness levels within a cell population are constantly monitored, ensuring the removal of suboptimal but otherwise viable cells (Merino et al., 2016). The elimination of potentially dangerous or abnormal cells based on their fitness status is known as cell competition. Recent findings prove cell competition is a broad biological process proposed to constitute a quality control mechanism against developmental malformations (de la Cova et al., 2004; Gibson and Perrimon, 2005; Moreno et al., 2002), tumorigenesis (Alexander et al., 2004; Hogan et al., 2009; Kajita and Fujita, 2015; Martins et al., 2014; Menéndez et al., 2010) and ageing (Merino et al., 2015). On the other hand, cell competition machinery may be subverted by pre-cancerous cells to acquire a super-fit status, enabling them to expand, kill and invade surrounding wild-type tissue with a lower fitness status (Eichenlaub et al., 2016; Levayer et al., 2015; Moreno and Basler, 2004; Suijkerbuijk et al., 2016). However up to date, cell competition was not yet investigated during the course of ageing-associated disorders, particularly in neurodegenerative diseases.

In *Drosophila* the fitness status is translated at the cellular level by ‘fitness fingerprints’, which are encoded by distinct isoforms of the Flower protein located at the extracellular membrane (Petrova et al., 2012; Rhiner et al., 2010; Yao et al., 2009). Flower is a conserved protein with three isoforms in *Drosophila* that differ solely at the extracellular C-terminus: Flower^ubi^ is expressed ubiquitously, while Flower^LoseB^ and Flower^LoseA^ are upregulated in suboptimal cells. The display of loser isoforms in a subset of cells is sufficient to target them for elimination by apoptosis, which is dependent on the transcription of the fitness checkpoint gene *azot* (Merino et al., 2015). Azot is an EF-hand calcium binding protein dedicated exclusively to cell competition-related apoptosis that integrates upstream relative fitness levels and targets suboptimal cells for death and subsequent engulfment by hemocytes (Portela et al., 2010, Casas-Tintó et al., 2015; Lolo et al., 2012). Mounting evidence demonstrates cell competition is a conserved process ranging from *Drosophila* to mammals that can also occur in post-mitotic cells and differentiated adult tissue such as epithelia or the neural system (Kolahgar et al., 2015; Tamori and Deng, 2013). The cell competition mediators *flower* and *azot*, for example, have been found to mediate elimination of injured or misconnected neurons (Merino et al., 2013; Moreno et al., 2015). The ‘flower code’ is cell type specific, since in the nervous system only Flower^LoseB^, and not Flower^LoseA^, is expressed in suboptimal neurons (Merino et al., 2013).

Neuronal loss is a key symptom of Alzheimer’s disease (AD), the most prevalent neurodegenerative disorder. AD is a slow progressive disease characterized by initial subtle memory problems that deteriorate to severe cognitive impairment, behavioural changes and difficulty to walk. The main pathological hallmarks of AD are brain deposition of extracellular amyloid-plaques and intracellular fibrils of hyperphosphorylated tau, exacerbated inflammation and finally neuronal damage and death (Braak and Braak, 1991). According to the amyloid cascade hypothesis, amyloid-β-related toxicity is considered the primary cause of the disease but the mechanisms mediating amyloid-induced neurodegeneration and cognitive decline are not fully elucidated (Ashe and Zahs, 2010; Huang and Mucke, 2012; Karran and De Strooper, 2016; Soldano and Hassan, 2014).

*Post-mortem* brain sections and structural MRI in AD patients show cerebral atrophy in regions involved in memory processing, such as the cortex and the hippocampus (Ossenkoppele et al., 2015; Seab et al., 1988). These findings suggest that the subpopulations of neurons primarily affected by AD, including the enthorhinal cortex and the hippocampal CA1 projection neurons, may be more vulnerable to cellular stress responses elicited by misfolded amyloid (Gómez-Isla et al., 1996; Saxena and Caroni, 2011; Wakabayashi et al., 1994). Although central to human pathology, mechanisms of neuronal loss have been understudied *in vivo* as AD mouse models do not recapitulate this aspect, showing little neuronal death (Ashe and Zahs, 2010; Karran et al., 2011).

Here we sought to analyze a potential role of fitness-based neuronal elimination in the context of AD onset and progression in *a Drosophila* model where human amyloid-β is induced in the adult fly brain. We found a physiological mechanism that identifies and purges less fit neurons, delaying cognitive decline and motor disability.

## Results

### Expression of amyloid-β42 in the *Drosophila* nervous system affects neuronal fitness

First, we tested whether neurons transit through a stage of reduced fitness when overexpressing Aβ42 (**Fig. 1A)**. We expressed a cassette containing two copies of the human amyloid-β42 (Aβ42) peptide fused to a signal peptide for secretion, under the control the *GMR-Gal4* driver, known to produce a strong degenerative phenotype in the *Drosophila* eye (**Fig. 1D)** (Casas-Tinto et al., 2011), onwards abbreviated as *GMR>Aβ42.* To monitor cell fitness markers in the optic lobe, where *GMR-Gal4* is expressed, we devised a sensitive reporter to detect Flower^LoseB^, by *knocking-in* a *flower^LoseB^::mCherry* tagged construct in the endogenous *flower* locus (**Fig. 1B)**. Flower^LoseB^*::mCherry* (indicator of low fitness) was strongly upregulated in the adult optic lobe of *GMR>Aβ42* flies but not in the *GMR>lacZ* control **(Fig. 1D,F)**.

**Figure 1.**
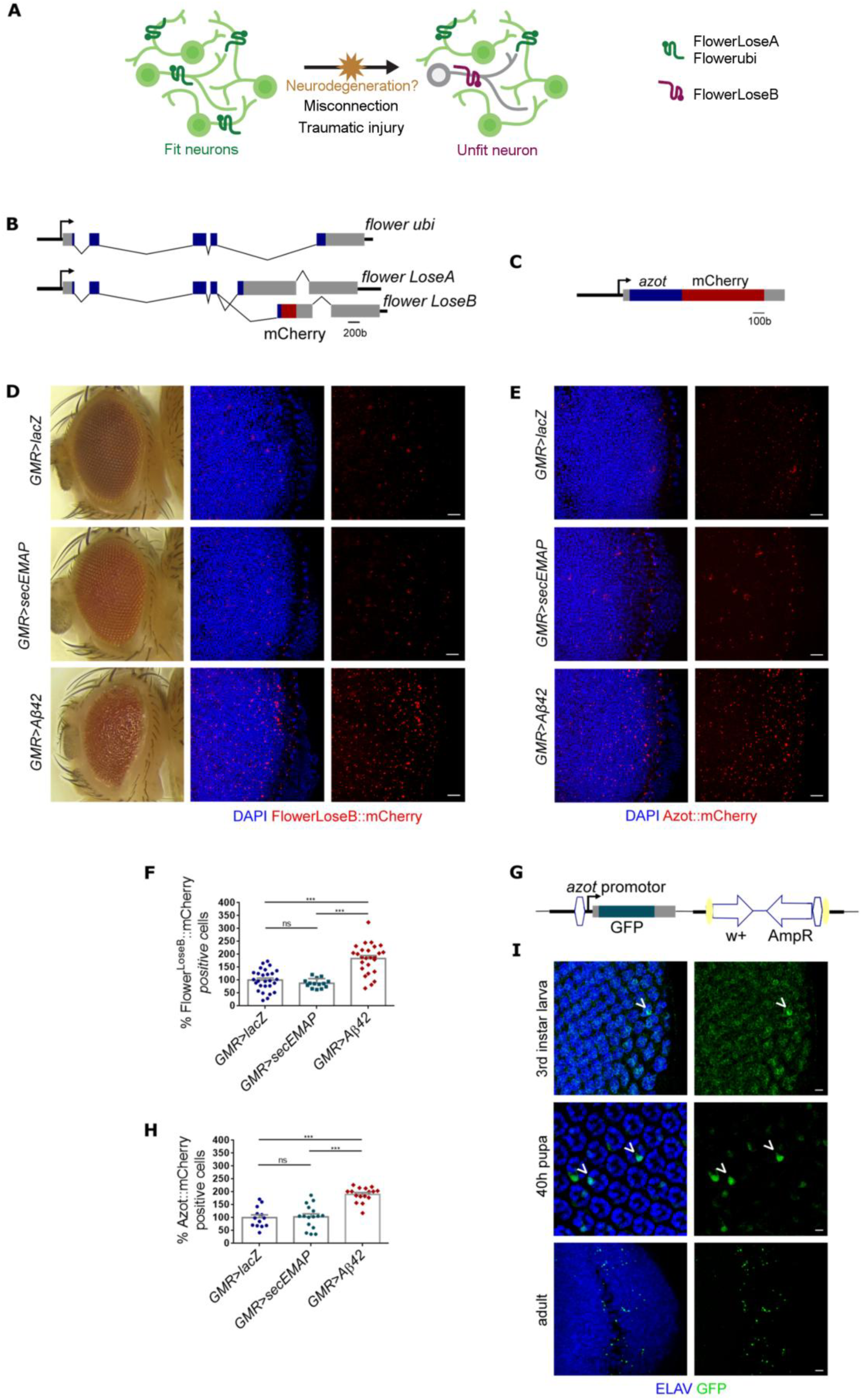
Expression of Aβ42 in the *Drosophila* nervous system generates suboptimal neurons that upregulate *flower^LoseB^* and *azot*. **(A)** Schematic illustrating neuronal fitness comparison. Neurons express ubiquitously Flower^ubi^ or Flower^LoseA^ at their cellular membrane, but external insults such as a traumatic injury or failing to establish connections during development can decrease the fitness status of one neuron (or one subpopulation of neurons) that upregulate Flower^LoseB^. We hypothesize that during the course of neurodegeneration induced by Amyloid-β, neurons enter a period of suboptimality, characterized by a low fitness status and upregulation of the Flower^LoseB^ isoform. (**B**) Representation of the *flower^LoseB^::mCherry* reporter. Each *flower* isoform has a different last exon. Based on this particularity, we generated a reporter specific for *flower^LoseB^* by introducing the mCherry sequence at the end of the exon specific for this isoform (exon 6). Blue rectangles are exons. The 5′ and 3′ untranslated regions are shown in grey. The red box shows the localization of the mCherry tag (not at scale). (**C**) Schematic of the *azot::mCherry* reporter that was obtained by fusion PCR. This construct includes 2430bp of the *azot* promoter region, the *azot* exon plus 175 bp of the 3’ end fused to mCherry (in red). Azot coding region is in blue and untranslated regions are represented in grey. **(D)** The *flower^LoseB^::mCherry* reporter (red) is strongly upregulated in the optic lobe of *GMR>*Aβ42 adults but not in the optic lobe of *GMR>lacZ or GMR>secEMAP* controls of the same age; the nuclear marker DAPI is shown in blue. Scale bar: 10 µm. The eye of *GMR>*Aβ42 flies shows a strong degenerative phenotype. **(E)** *azot::mCherry* reporter (red) expressed in the optic lobe of adult flies in the presence of *GMR*-driven *lacZ*, *secEMAP* or *Aβ42*; DAPI is shown in blue. Scale bar: 10µm. **(F)** Quantification of the percentage of Flower^LoseB^::mCherry positive cells in the optic lobes of the indicated genotypes. The number of Flower^LoseB^::mCherry positive cells detected for the *GMR>lacZ* control group was assumed as 100%. **(G)** Schematic of the modified *azot{KO; GFP}* locus. This transgenic line was generated by integration of a *knock-in* construct containing the GFP sequence under the control of the *azot* endogenous promoter, into the *azot knock-out* locus. The 5′ and 3′ untranslated regions of the *azot* gene are shown in grey. The vector backbone was conserved in the *knock-in* line (*w*+, *AmpR*). The yellow ellipses are *loxP* sites and the white hexagons are *attL* regions. **(H)** Quantification of the percentage of Azot::mCherry positive cells in the optic lobe of the indicated genotypes. The number of Azot::mCherry positive cells for the *GMR>lacZ* control group was assumed as 100%. **(I)** Eye imaginal discs of third instar larva, retina of 40h pupa and adult optic lobes of *GMR>Aβ42* adults showing immunolabelling for the nuclear marker ELAV (blue) and endogenous GFP signal produced from *azot{KO; GFP}* (green). Arrow heads point to co-localization between ELAV and GFP. Scale bar: 5 µm. Error bars show standard error mean. Ns: no significant. ***P value<0,001. All genotypes are heterozygous. See also Figure S1 and Figure S2.

To control if the secretion of a peptide *per se* is sufficient to downregulate fitness levels and induce *flower^LoseB^*, we expressed the secreted form of a small peptide, EMAP (17kd), under the control of *GMR-Gal4*. Secreted EMAP is a chemotactic clue that attracts haemocytes to sites of cell competition (Casas-Tintó et al., 2015). We confirmed that secretion of EMAP alone does not upregulate Flower^LoseB^*::mCherry* in the optic lobe, indicating that secretion of an innocuous peptide is not sufficient to decrease the fitness levels of neurons **(Fig. 1D,F)**.

The Flower^LoseB^ isoform was particularly upregulated in neurons of the optic lobes as detected with the neuronal marker Elav **(Fig. S1A)**. Accordingly, Flower^LoseB^ expression did not co-localize with cells expressing the glial marker Repo **(Fig. S1B)**.

We then tested activation of another marker of low fitness *azot*, which is transcribed in cells destined to die based on previous fitness comparison (Merino et al., 2015). In order to visualize *azot* expression, we generated 1) *azot::mCherry* transgenic flies, which carry an extra copy of *azot* fused to *mCherry,* inserted in another chromosome **(Fig. 1C)** and 2) *azot{KO;GFP}* flies, wherein *GFP* was placed in the endogenous locus of the previously *knocked-out* (KO) *azot* gene **(Fig. 1G)**. With both lines, we found that *azot* was not only upregulated in the optic lobes of *GMR>Aβ42* adult flies but was already activated in neurons at previous developmental stages, including the eye discs of the larva and retinas of mid-pupa **(Fig. 1E,H,I)**.

### Expression of misfolding-prone toxic peptides linked to Huntington’s, but not to Parkinson, triggers neuronal competition

To further investigate neuronal fitness comparison in other types of neurodegenerative diseases, we turned to published human trangenes reported to induce degenerative phenotypes in the fly: HttQ128 and α-SynA30P. HttQ128 is a pathogenic form of the human *huntingtin* gene that encodes an expanded repeat of 128 poly-glutamines, causing reduction of viability, retinal death and abnormal motor behaviour in *Drosophila* (Lee et al., 2004). α-SynA30P is a mutant allele linked to familial Parkinson disease that originates premature loss of dopaminergic neurons, formation of brain inclusions similar to Lewis bodies and decrease of climbing ability when expressed in flies (Feany and Bender, 2000; Song et al., 2017). HttQ0 and α-SynWT, which carry a non-pathogenic form of *huntingtin* and the wild-type allele of *α-synuclein*, respectively, served as controls.

For this experiment we employed a previously published translational reporter, in which Flower^UBI^, Flower^LoseA^ and Flower^LoseB^ are tagged with a specific fluorescent protein: YFP, GFP and RFP, respectively (Yao et al., 2009). We discovered that expression of HttQ128 from the *GMR* driver induces augmented levels of Flower^LoseB^ in the adult brain, contrary to the non-pathogenic form, HttQ0 **(Fig. 2A,B)**. Surprisingly, levels of Flower^LoseB^ did not change with ectopic expression of the Parkinson-related peptides, α-SynA30P and α-SynWT **(Fig. 2E,F)**. The same results were obtained using the Flower^LoseB^::mCherry reporter to detect changes on cell fitness upon expression of these toxic peptides in the eye imaginal disc of the larva **(Fig. S2A-C)**.

**Figure 2.**
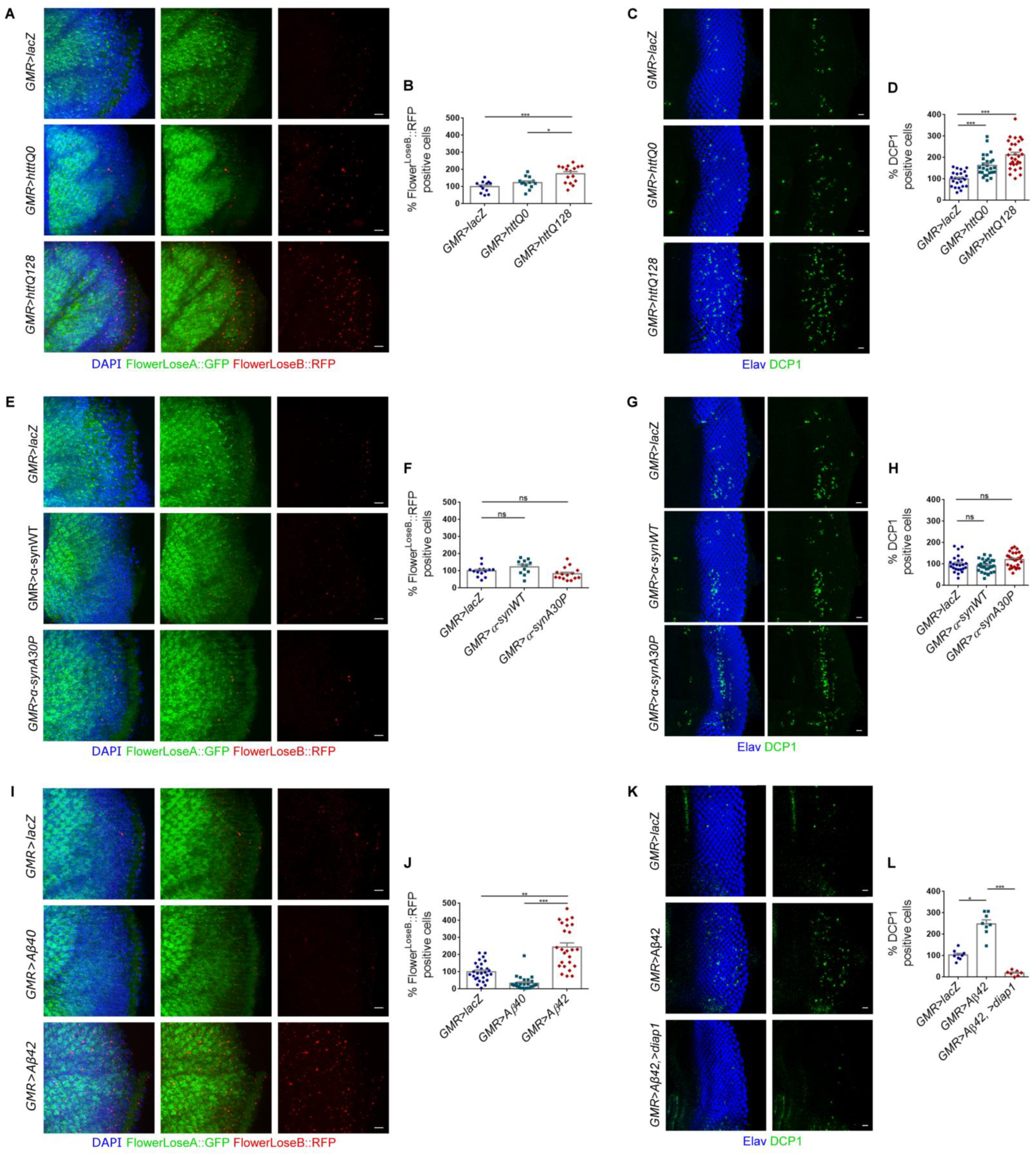
Ectopic expression of HttQ128, but not α-SynA30P, induces an upregulation of the Flower^LoseB^ isoform. (**A,B**) Expression of the *flower^ubi^::YFP, flower^LoseA^::GFP, flower^LoseB^::RFP* reporter in the optic lobe of adult flies and quantification of the RFP positive cells (red) for the following genotypes: *GMR>lacZ*, *GMR>httQ0* and *GMR>httQ128*. DAPI is shown in blue and GFP in green. Scale bar: 10 µm. (**C,D**) Quantification of the number of apoptotic cells in the eye discs of third instar larva expressing *UAS-lacZ*, *UAS-httQ0* or *UAS-httQ128* under the control of *GMR-Gal4* driver and representative figures for each genotype. Apoptotic cells are marked by DCP1 in green and nuclei are shown in blue. Scale bar: 10 µm. (**E,F**) Expression of the *flower^ubi^::YFP, flower^LoseA^::GFP, flower^LoseB^::RFP* reporter in the optic lobe of adult flies and quantification of RFP positive cells (red) for the following genotypes: *GMR>lacZ*, *GMR>α-synWT* and *GMR>α-synA30P*. DAPI is shown in blue and GFP in green. Scale bar: 10 µm. (**G,H**) Quantification of the number of apoptotic cells in the eye discs of third instar larva expressing *UAS-*l*acZ*, *UAS-*α-*synWT* or *UAS-*α-*synA30P* under the control of *GMR-Gal4* and representative figures for each genotype. Apoptotic cells are marked by DCP1 in green and nuclei are represented in blue. Scale bar: 10 µm. (**I,J**) Expression of the *flower^ubi^::YFP, flower^LoseA^::GFP, flower^LoseB^::RFP* reporter in the optic lobe of adult flies and quantification of RFP positive cells (red) for the following genotypes: *GMR>lacZ*, *GMR>Aβ40* and *GMR>Aβ42*. DAPI is shown in blue and GFP in green. Scale bar: 10 µm. (**K,L**) Quantification of the number of apoptotic cells in the eye discs of third instar larva expressing *UAS-lacZ*, *UAS-Aβ42* or *UAS-Aβ42/UAS-diap1* under the control of *GMR-Gal4* and representative figures for each genotype. Apoptotic cells are marked by DCP1 in green and nuclei are represented in blue. Scale bar: 10 µm Error bars indicate S.E.M. *P value<0,05. **P value<0,01. ***P value<0,001. The number of positive cells for the genotype *GMR>lacZ* was assumed as 100% for normalization. See also Figure S2.

Although both HttQ128 and α-SynA30P induce neurodegeneration by accumulation of protein aggregates in *Drosophila* models, our results indicate that only HttQ128 triggers neuronal competition. This result may be explained by the difference in toxicity levels imposed to the tissue by each of these transgenes. We observed that HttQ128 expression leads to increased cell death in a larval epithelium (eye disc) in opposition to the α-SynA30P transgene, which did not lead to significantly increased apoptosis under the same conditions **(Fig. 2C,D,G,H)**.

Accumulation of amyloid peptides in the brain is thought to be the first step in Alzheimer’s pathogenesis. While Aβ42 is the main component of amyloid plaques found in human patients, Aβ40 is a shorter isoform of the human amyloid peptide, which is less amyloidogenic. Aβ40 does not deposit as soluble oligomeric aggregates in *Drosophila* and does not cause neurodegeneration *in vivo* (Iijima et al., 2004; Speretta et al., 2012). When overexpressed in the *Drosophila* brain using *GMR-Gal4*, Aβ40 did not alter the levels of the Flower^LoserB^ reporter **(Fig. 2I-L)**. We decided to focus on Aβ42-associated toxicity from now on because the levels of *azot* and *flower* were most affected by this peptide.

### Aβ42-producing clones are eliminated over time from a neuronal epithelium

To determine if Aβ42 induces cell elimination when expressed in clones, we induced its expression by heat-shock in clones marked by GFP in the neuro-epithelium of the eye disc. We registered that Aβ42-producing clones are progressively excluded from the tissue and detected a higher proportion of dying cells marked by DCP1 inside these clones **(Fig. 3A,D,E,F)**. Flower^LoseB^::mCherry and Azot::mCherry were majorly detected inside Aβ42-producing clones, but some signal was also present outside of clones borders **(Fig. 3B,C,E,F)**. We found that Aβ42 diffused largely out of clone borders and accumulated at the basal side of the eye discs, explaining non-autonomous induction of *flower* and *azot* (**Fig.S2D)**. Flower^LoseB^::mCherry and Azot::mCherry were not detected in control clones expressing an innocuous transgene **(Fig.3C).** As expected the cleaved form of *Drosophila* Caspase Protein 1 (DCP1) co-localized with Flower^LoseB^::mCherry, showing that unfit cells affected by Aβ42 were undergoing apoptosis (**Fig. 3E, Fig.S2E)**.

**Figure 3.**
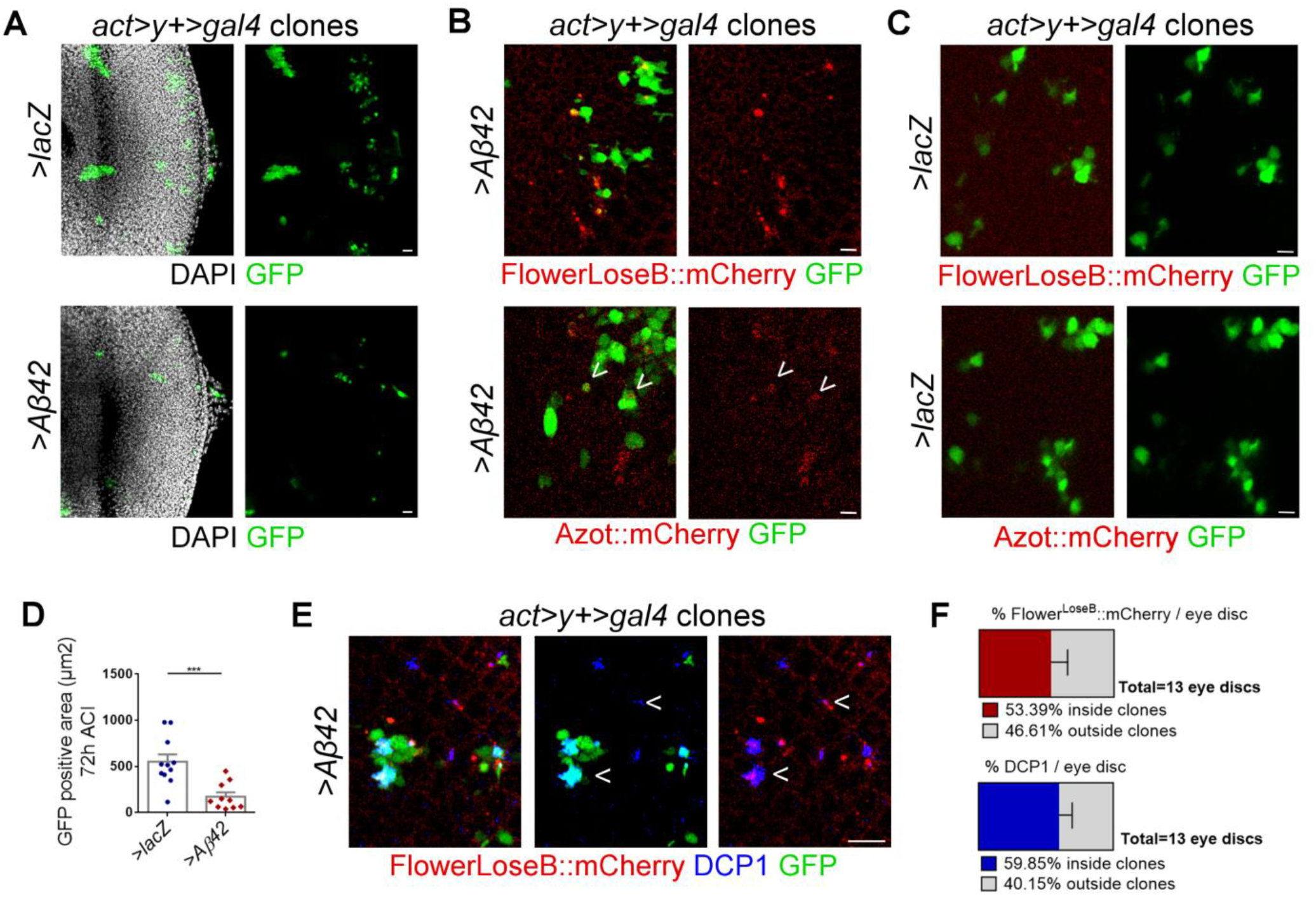
Aβ42-producing clones are eliminated over time from a neuroepithelium. **(A)** Clones induced by heat-shock of the flip-out cassette *act>y+>gal4,UAS-GFP* in eye imaginal discs of third instar larva. Clones are marked by GFP (green) and express *UAS-lacZ* or *UAS-Aβ42,* 72h after clone induction (ACI). DAPI is in white. Scale bar: 20 µm. **(B)** Expression of the Flower^loseB^::mCherry reporter (red) 48h ACI or of the Azot::mCherry reporter (red) 72h ACI in *Aβ42*-overexpressing clones (green). Arrow heads indicate co-localization. Scale bar is 5µm. **(C)** Clones (green) induced by heat-shock of the flip-out cassette *act>y+>gal4* driving *UAS-lacZ* in the eye imaginal disc of third instar larva. No expression of the Flower^loseB^::mCherry reporter (red) or the Azot-mCherry reporter (red) is detected 48h ACI or 72h ACI, respectively. Scale bar: 5µm. **(D)** Quantification of GFP-positive area in *flip-out* clones overexpressing *lacZ or Aβ42* 72h ACI. **(E)** Detection of cleaved DCP1 (blue) and Flower^LoseB^::mCherry (red) expression in *act>y+>gal4,UAS-GFP* clones (green) driving Aβ42 secretion, 48h ACI. Arrows indicate co-localization. Scale bar: 10µm. **(E)** Quantification of the percentage of Flower^LoseB^::mCherry signal or DCP1 positive signal localizing inside or outside *act>y+>gal4* clones expressing Aβ42 48h ACI. See also Figure S2.

### *flower* and *azot* are necessary for amyloid-β42 induced neuronal death

Next, we analysed Aβ42-associated toxicity and neuronal loss in the adult brain. *GMR-*driven Aβ42 induces cell death in the optic lobe over time, eliciting a 2.8-fold increase in the number of positive cells for activated DCP1, which co-localized with the neuronal marker ELAV, compared to control flies at two weeks after eclosion **(Fig. 4B,C)**.

**Figure 4.**
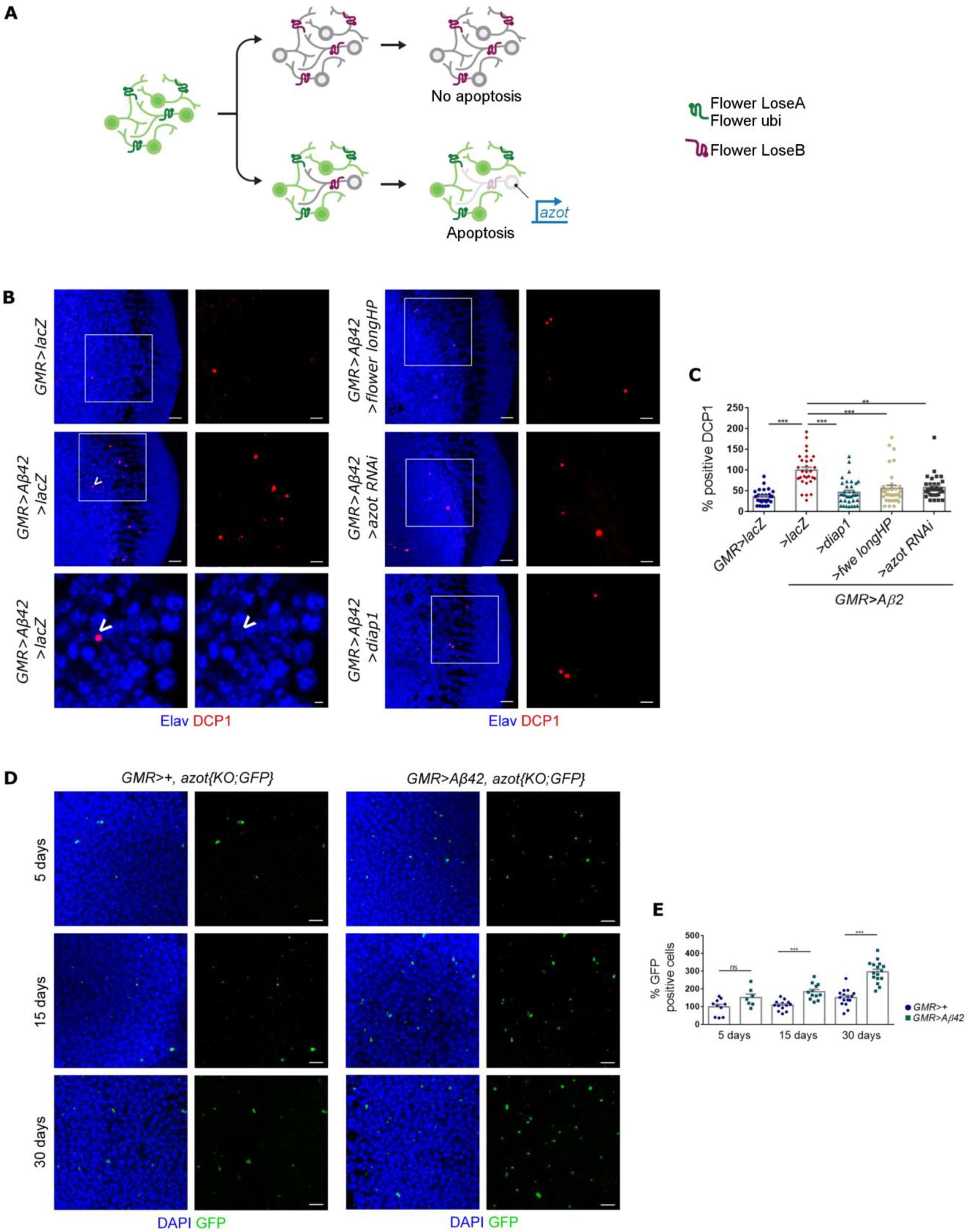
*flower* and *azot* are necessary for amyloid-β42 induced neuronal death. **(A)** Neurons compute relative levels of Flower^LoseB^ to determine its fate. Neurons expressing LoseB and surrounded by neurons with equal levels of LoseB do not activate *azot* transcription. If the neuron cannot cope with an insult and exhibits persistently higher relative levels of LoseB compared to neighboring neurons, it will fail to pass a fitness checkpoint mediated by the *azot* gene and will be purged from the tissue. Increased copies of *azot* enhance fitness-based cell elimination. **(B)** Optic lobes of two weeks old adults showing apoptotic neurons labeled by DCP1 (*Drosophila* Caspase Protein1) in red and ELAV in blue. Scale bar: 10 µm or 5 µm in the insets. At the bottom on the left panel is shown a dying neuron (arrow) from a single plane of the confocal projection displayed above, representative of the *GMR>Aβ42 / >lacZ* genotype (scale bar is 2µm). **(C)** Quantification of positive DCP1, assuming the levels of apoptosis in *GMR>Aβ42 / >lacZ* as 100%. **(D)** Tracing of suboptimal cells (green) through ageing using the *azot{KO; GFP}* reporter in *GMR>+ flies* or *GMR>Aβ42 flies* at 5,15 or 30 days of life. DAPI marks cell nuclei (blue). Scale bar: 5 µm. **(E)** Quantification of GFP-positive cells per optic lobe in *GMR>+* flies or *GMR>Aβ42* flies at 5,15 or 30 days of life. Error bars show S.E.M. Ns: no significant. **P value<0,01. ***P value<0,001. All genotypes are heterozygous. See also Figure S3.

The presence of Flower^LoserB^ isoforms at the cell membrane of a particular neuron does not imply that the cell will die (Merino et al., 2013; Moreno et al., 2015; Rhiner et al., 2010) **(Fig. 4A)**. Only if relative fitness differences with neighbouring neurons persist, cell death is initiated (Levayer et al., 2015; Merino et al., 2013; Rhiner et al., 2010), which requires downstream transcriptional activation of *azot* (Merino et al., 2015) **(Fig. 4A)**.

To check if neuronal fitness comparisons mediate Aβ42-induced death, we modulated *flower* and *azot* genetic dosages. We found that suppressing relative differences of Flower^LoseB^ levels among cells by silencing *LoseA/B* isoforms via a long hairpin (Merino et al., 2013), is sufficient to induce a strong decrease in total apoptosis detected upon Aβ42-expression in the adult brain, bringing it back to almost *wild-type* levels (**Fig. 4B,C**). Silencing *azot* with *RNAi* also reduced the cell death observed in the presence of Aβ42 alone, bringing it back to almost *wild-type* levels **(Fig. 4B,C**).

As a positive control for inhibition of apoptosis we used *UAS-dIAP1* (*>dIAP1*), an antagonist of the apoptotic pathway in *Drosophila* (Hay et al., 1995). Total cell death in *GMR>Aβ42* adult optic lobes was suppressed by over-expression of dIAP1 (*Drosophila inhibitor of apoptosis1*) **(Fig. 4B,C)**. We confirmed that part of apoptotic cells marked by DCP1 co-localize with Azot::mCherry in a *GMR>Aβ42* background **(Fig. S1C)**. These results were further supported by a similar experiment conducted in the eye imaginal disc of the larva **(Fig. S3A,B)**, showing that Aβ42-associated cell death is mediated by *flower* and *azot.* Moreover we found that the cell fitness-based neuronal elimination induced by neurodegeneration is not specific to *Aβ42*, and also occurs in the case of HttQ128-associated degeneration **(Fig. S3C,D)**.

In order to study how neuronal fitness is affected over time we monitored cumulative *azot* expression during ageing of *GMR>Aβ42* brains with the reporter line *azot{KO;GFP}*. This reporter allows visualization of impaired cells (GFP+) that activate the *azot* promoter (Merino et al., 2015). Using this tool, we observed that GFP signal is only detected in Flower^LoseB^ positive cells **(Fig. S1D)**. Optic lobes overexpressing Aβ42 accumulate GFP-positive cells at an increased rate compared to control brains of identical ages lacking Aβ42 (53%, 70% and 96% increase over the *wild-type* of the same age at 5, 15 and 30 days, respectively) **(Fig. 4D,E)**. Altogether, this shows that Aβ42 expression leads to a progressive generation of neurons which will be targeted to death via *azot* **(Fig. 4A).**

### Suppression of *azot*-dependent removal of Aβ42-damaged neurons aggravates accumulation of degenerative vacuoles and decreases lifespan

Next, we asked what are the consequences of blocking fitness-based elimination of Aβ42-damaged cells for ageing, locomotion and cognition.

First we established a model where expression of *Aβ42* is restricted to adult neurons: To this end, we generated flies containing the *Aβ42*-cassette under the control of the inducible promoter *elav-GeneSwitch (elavGS) (***Fig. 5A)** (Poirier et al., 2008; Roman et al., 2001). We detected Aβ42 accumulation in the optic lobes and mushroom body calyx of *elavGS>Aβ42* adult flies fed for 5 days on the *GeneSwitch*-activator RU486 but not in the brain of uninduced flies (**Fig. 5B, Fig. S4A)**. Aβ42 aggregates stained positive for aggresome markers **(Fig. S4G)**, confirming the amyloidogenic nature of human Aβ42 when secreted by *Drosophila* neurons.

**Figure 5.**
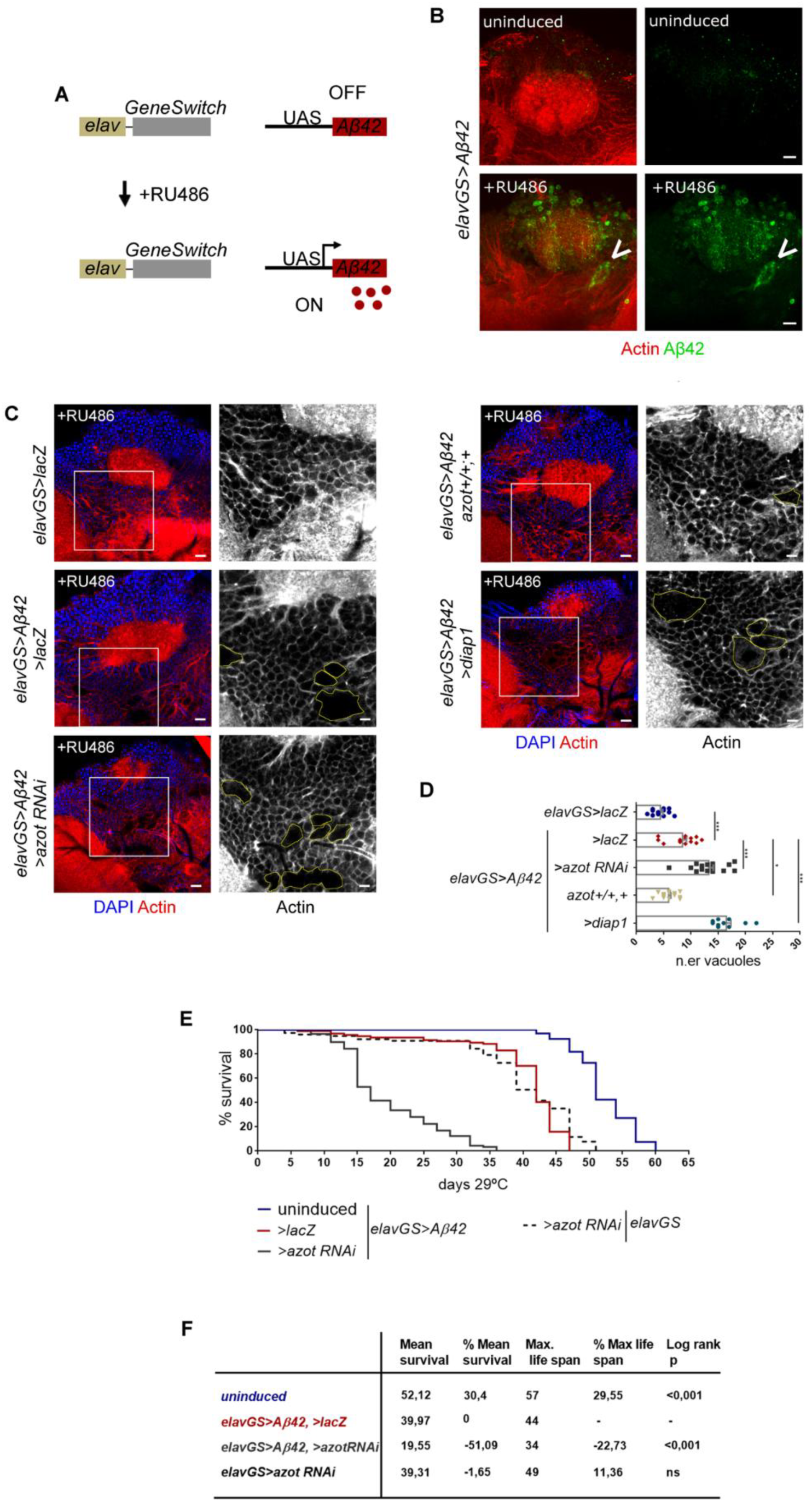
Suppression of fitness-based neuronal selection decreases lifespan in a *Drosophila* model of AD. **(A)** The *GeneSwitch* system relies on a chimerical Gal4 containing an esteroid receptor domain that only becomes active upon binding the synthetic progesterone analog RU486. This system allows us to induce the conditional expression of the human Aβ42 peptide in the adult brain of *elav-GeneSwitch (elavGS)* transgenic flies upon addition of RU486 to the food. **(B)** Aβ42 (green) expression is detected in the brain of RU486-induced *elavGS>Aβ42* flies but not in the brain of uninduced flies. Aβ42 can form insoluble aggregates with large size (arrow). A posterior view of the brain next to the mushroom body calyx is shown. Actin is in red. Scale bar: 5 µm. **(C)** Posterior view of the brain, showing the surrounding region of the mushroom body calyx. Nuclei are marked by DAPI (blue) and actin cytoskeleton by phalloidin (red). Degenerative vacuoles are surrounded by a yellow line in grayscale insets. All genotypes were treated with RU486. Scale bar: 10µm in color pictures or 5µm in grayscale insets. **(D)** Mean number of vacuoles located at a 10µm-deep plan in two-weeks old brains of the indicated genotypes. Data are represented as mean±SEM. ***P value<0,001, **P value<0,01, *P value<0,05. **(E,F)** Lifespan curve and table depicting survival analysis for heterozygous females of the following genotypes: uninduced *elavGS>*Aβ42 / >*lacZ*; induced elavGS*>Aβ42 / >lacZ;* induced *elavGS>Aβ42 / > azot RNAi and* induced *elavGS>azot RNAi.* See also Figure S4 and Figure S5.

Apoptosis was increased in the optic lobes of 10 days-old adults raised on RU486 **(Fig. S4B)** compared to uninduced flies. TUNEL-positive cells co-localized with ELAV, indicating that Aβ42 caused neuronal death **(Fig. S4B).** RU486 did not cause apoptosis on its own (**Fig. S4C,D**). Increased cell death was not accompanied by elevated proliferation, consistent with the fact that little regeneration occurs in uninjured adult brains (Fernández-Hernández et al., 2013) (**Fig. S4E,F**).

Brains of induced *elavGS>Aβ42* flies showed hallmarks of neurodegeneration, such as increased number of degenerative vacuoles **(Fig. 5C,D).** In induced *elavGS>Aβ42* flies, the total number of vacuoles was the double of *elavGS>lacZ* control flies of the same age (**Fig. 5C,D)**. Interestingly, *azot knock-down* in induced *elavGS>Aβ42* further aggravated brain degeneration and caused *a* 57% increased in the total number of neurodegenerative vacuoles **(Fig. 5C,D)** Conversely, when induced *elavGS>Aβ42* flies were provided with a third functional copy of *azot*, which is known to accelerate the elimination of unfit cells (Merino et al., 2015), brain architecture was restored and the number of vacuoles dropped 30% (**Fig. 5C,D**). Finally, we suppressed apoptosis by overexpressing dIAP1 together with Aβ42 in adult neurons and observed that brains deteriorated faster than in induced *elavGS>Aβ42* flies alone (**Fig. 5C,D).**

To rescue brain morphology, we made use of the *azot{KO;hid}* transgenic line, which contains the coding sequence of the pro-apoptotic gene *hid* inserted in the *azot KO* locus, leading to *hid* transcription under the control of *azot* endogenous enhancer sequences (Merino et al., 2015). The total size of vacuoles in the brain of induced *elavGS>Aβ42/ azot{KO;hid}* flies, which lack Azot protein but still eliminate unfit cells via transcription of *hid*, was significantly decreased at two weeks time, proving *azot* has a role mainly dedicated to apoptosis regulation in the course of neurodegeneration (**Fig. S5A,B)**.

The observation that suppression of apoptosis led to accelerated vacuole formation in the brain, made us speculate that cells may be undergoing alternative forms of cell death, such as necrosis. To test this hypothesis, we followed a protocol using propidium iodide (PI) that can penetrate compromised membranes of necrotic cells (Liu et al., 2014; Yang et al., 2013). We detected an increased number of cells permeable to PI in the brains of *elavGS>Aβ42* flies two weeks after induction, comparing to non-induced flies, indicating that Aβ42 can trigger necrosis in the brain (**Fig. S5C**). However, blocking apoptosis either by overexpression of dIAP1 or *knock-out* of *azot* did not lead to a further increase in the levels of necrosis, making necrosis an unlikely cause of accelerated vacuole formation in these genotypes **(Fig. S5C).**

When analysing life expectancy of *elavGS>Aβ42* flies, we found that secreted Aβ42 is detrimental for longevity (**Fig. 5E,F)**: induced flies lived on average 40 days versus 52 days for uninduced flies. Life expectancy of induced *elavGS>Aβ42* flies dropped to 20days when *azot* expression was silenced by RNAi (representing a 51% decrease in mean survival comparing with *elavGS>Aβ42* flies carrying a wild-type dose of *azot*) (**Fig. 5E,F)**. It was not possible to determine a clear effect of *diap1-*overexpression on longevity *per se* or in combination with Aβ42 (data not shown). Apoptosis is involved in many biological processes with potentially opposing consequences for lifespan.

### An extra copy of *azot* is sufficient to restore motor coordination and improve long-term memory formation

Furthermore, we studied the consequences of fitness-based neuronal-culling on walking behaviour. Using tracking software (Colomb et al., 2012) we extracted several behavioural parameters from 5 min walking sessions of individual flies (**Fig. 6A**). *elavGS>Aβ42* flies induced for two weeks on RU486 showed decreased activity time, shorter walks and ataxia (**Fig 6B**), compared to uninduced *elavGS>Aβ42* flies (**Supplementary Videos 1,2)**. *azot* RNAi significantly exacerbated behavioural and locomotor dysfunctions caused by Aβ42 alone (**Fig. 6A,B**). On the contrary, an extra copy of *azot* was sufficient to completely restore the behavioural defects observed in *elavGS>Aβ42* flies, including lengths of walks, activity time and ataxia (**Fig. 6A,B, Supplementary Video 3).** Finally, blocking apoptosis with *UAS-diap1* in *elavGS>Aβ42* individuals, further compromised walking performance (**Fig. 6A,B)**, whereas dIAP1-overexpression alone did not result in impaired locomotion (**Fig. S6**).

**Figure 6.**
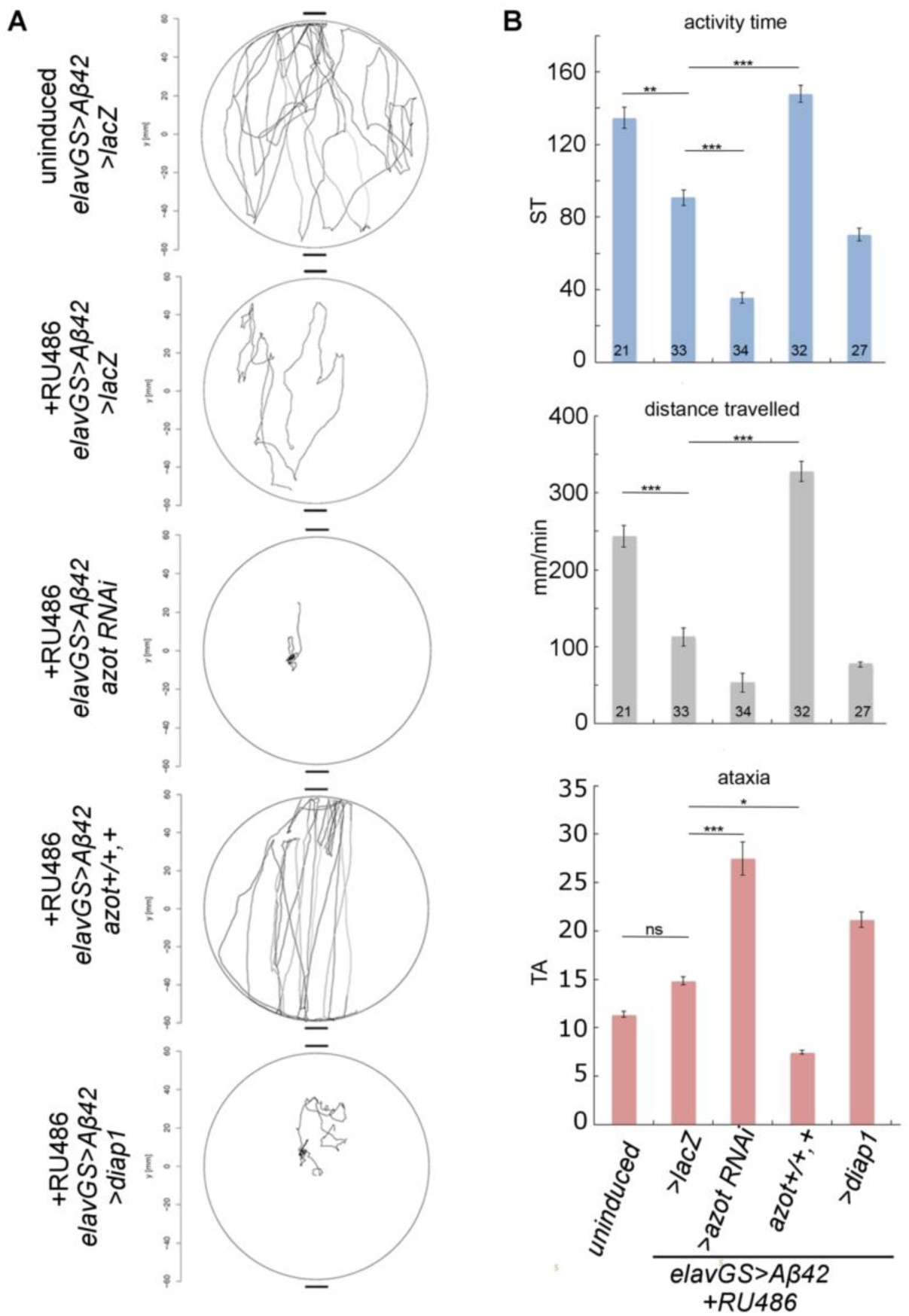
An extra copy of *azot* is sufficient to restore motor coordination in *AD* flies. **(A)** Individual walking trajectories (5min tracking) representative of each indicated genotype. **(B)** Graphs depict activity time in seconds (using a speed threshold, ST)(Colomb et al., 2012), distance walked in mm/min and median turning angle TA in degrees. Parameters were calculated from individual walks of two weeks-old heterozygous flies raised on RU486 for each genotype. Statistical significance based on ANOVA. All genotypes were compared to each other.

To assess long-term memory (LTM) formation we used courtship suppression assays (Keleman et al., 2007; Siegel and Hall, 1979). Courtship conditioning is a form of associative learning by which male flies have to recall that they were previously rejected by a mated/unreceptive female and reduce courtship activity when re-exposed **(Fig. 7A)**. Because prolonged *Aβ42* expression resulted in locomotion defects, we reduced *Aβ42* induction by one week to ensure that all naïve control males reached a courtship index of 0.6-0.8 (courting 60%-80% of the observation period) (**Fig. 7B**), normally seen in wild-type sham controls (Nichols et al., 2012). Presence of RU486 did not affect LTM formation of *elavGS* flies without the *Aβ42* transgene (data not shown). We measured a significant difference in courtship index between sham and trained males for all genotypes except for *elavGS>Aβ42, azotKO^-^/-* flies **(Fig 7B)**. One week-induced *elavGS>Aβ42* flies showed impaired LTM formation compared to uninduced flies **(Supplementary Video 4,5)**, which was strongly aggravated in the absence of *azot* (**Fig. 7B,C**) **(Supplementary Video 6).** Additional expression of *>diap* to block cell death had a detrimental effect on LTM (but not statistical significant) **(Fig 7C and Supplementary Video 7)**. Conversely, introduction of an extra copy of *azot,* which increases the efficiency of cell culling (Merino et al., 2015), was sufficient to restore robust LTM formation in A*β42*-expressing flies **(Fig 7C,D and Supplementary Video 8)**, resulting in a significant improvement of memory compared to *elavGS>Aβ42 / >lacZ* flies. These results underline that *azot*-mediated clearance of neurons is beneficial for motor and cognitive functions affected by adult onset *Aβ42-*expression. Moreover, introduction of a single extra copy of *azot* was sufficient to completely prevent *Aβ42*-induced motor and cognitive decline.

**Figure 7.**
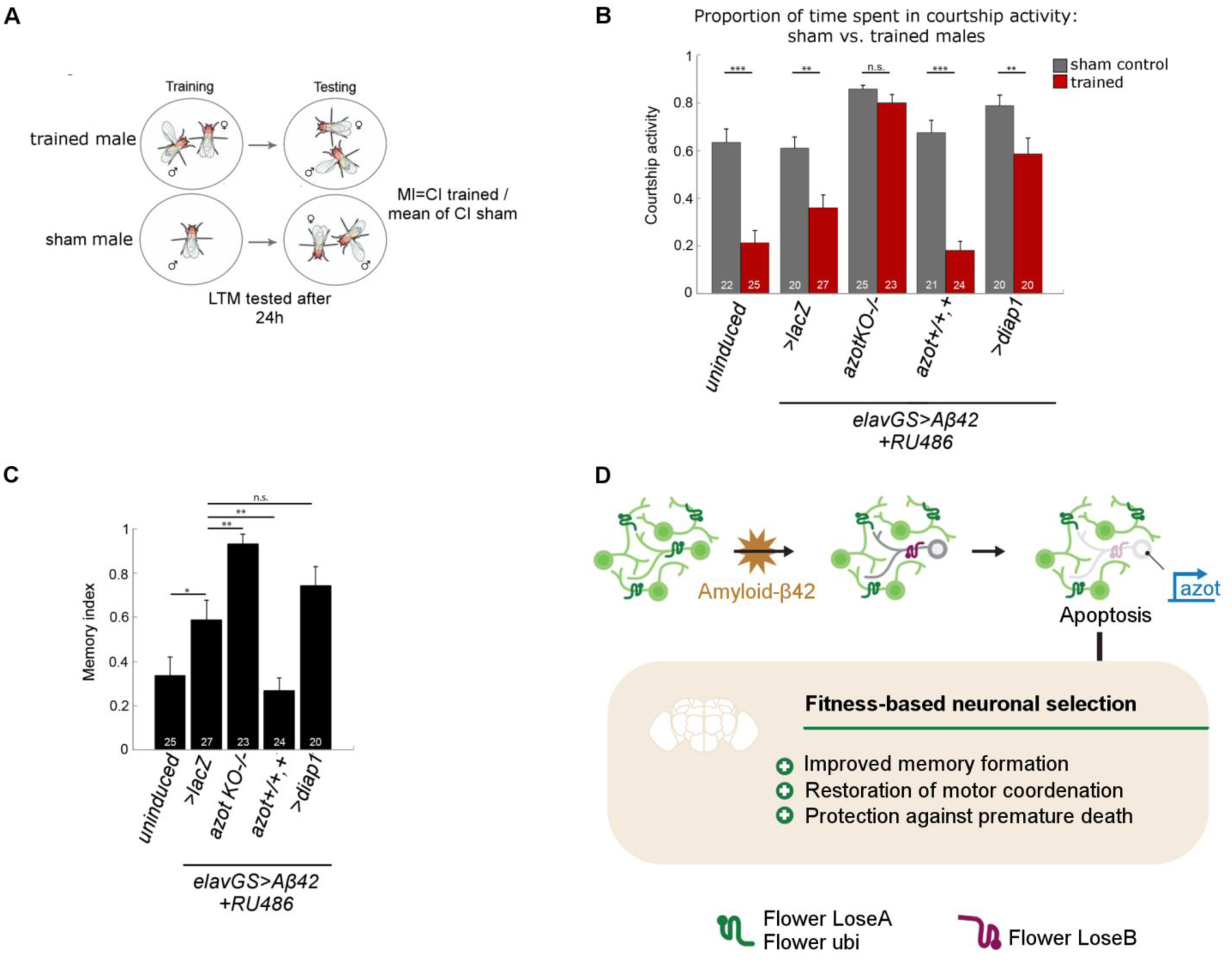
An extra copy of *azot* improves long-term memory formation in flies expressing Amyloid-β42. **(A)** Schematic depicting the courtship suppression assay to measure long-term memory. **(B)** Graph showing the courtship index of sham (grey bars) vs trained males (red bars) of the indicated genotypes. Statistical analysis based on Student’s T-test. **(C)** Graph showing the memory index of the indicated genotypes. Statistical analysis based on ANOVA **(D)** Amyloid-β42-induced neuronal death is mediated by fitness comparison encoded by *flower* and executed by *azot*. Removal of amyloid-β42-compromised neurons via cell competition has a strong net beneficial effect at the organismal level, protecting against motor decline, memory impairment and premature death. Error bars show S.E.M, numbers within the bars indicate the number of individuals tested. *** P value < 0.001, ** P value < 0.01, * P value < 0.05. See also Figure S6.

In flies, the mushroom body (MB) is important for learning and memory (Aso et al., 2014). To investigate if the above observed memory defects were caused by altered MB architecture, we revealed MB structure by anti-Fasciclin II (FasII) immunohistochemistry, which strongly labels the α and β lobes (Crittenden et al., 1998; Fushima and Tsujimura, 2007). We analyzed all genotypes after one week of Aβ42 induction when memory phenotypes were evident, but did not detect marked differences in MB structure. In particular, *elavGS>Aβ42, azotKO-/-* flies did not exhibit severe morphological defects despite the strong memory impairment **(Fig S6).** We observed a modest variability between individuals of the same genotype and depicted mild alterations in lobes of the MB, which were comparable among genotypes **(Fig S6)**. This result suggests that memory differences between genotypes are not a result of MB malformation but rather a consequence of a genetic interaction between Aβ42 and *azot*.

## Discussion

Here we found that expression of misfolding-prone toxic peptides linked to Alzheimer’s and Huntington’s diseases in *Drosophila* neural tissues affects neuronal fitness and triggers cell competition, leading to increased activation of the Flower^LoseB^ isoform and Azot. Our results demonstrate that fitness fingerprints are important physiological mediators of cell death occurring during the course of neurodegenerative diseases. However this mechanism is specific to certain neurodegeneration-causing peptides or particular stages of the disease, since it is not elicited by expression of Parkinson-related α-Synuclein, for instance. Interestingly, our results suggest that that the toxic effects of a given peptide correlate directly with the level of neuronal competition and death it induces.

Surprisingly we found that neuronal death had a beneficial effect against β-amyloid-dependent cognitive and motor decline. This finding challenges the commonly accepted idea that neuronal death should be detrimental at all stages of the disease progression. We found that most amyloid-induced neuronal apoptosis is beneficial and likely acts to remove damaged and/or dysfunctional neurons in an attempt to protect neural circuits form aberrant neuronal activation and impaired synaptic transmission.

One curious observation in our study is the fact that Aβ42 induces cell death both autonomously and non-autonomously in clones of the eye disc. Dying cells co-localize with Flower^LoseB^ reporter both inside and outside of GFP-marked clones. We observed that Aβ42 peptide is secreted to regions outside of clones and accumulates at the basal side of the eye disc. The neurons of the eye disc, which project their axons into the optic stalk through the basal side of the disc, are likely affected by the accumulation of this toxic peptide, explaining the induction of cell death outside of clones.

We detected that blocking apoptosis in amyloid-β42 expressing flies either by *azot* silencing or overexpression of dIAP1 increases the number of vacuoles in the brains of these flies. This seems to be a counterintuitive observation as one would expect that a reduction in apoptosis would result in less cells being lost and a reduction of neurodegenerative vacuoles. However this observation can be conciliated with our model: we suspect that less fit neurons have impaired dendritic growth and inhibit the expansion of neighbouring neurons. This inhibition would disappear once the unfit neuron is culled, allowing compensatory dendritic growth and neuropil extension.

The fact that introduction of a single extra copy of *azot* was sufficient to completely prevent *Aβ42*-induced motor and cognitive decline may suggest new venues for AD treatment that aim to support elimination of dysfuncional neurons at early stages of AD pathology. For example, in patients at early symptomatic stages when cognitive impairment is first detected, enhancing physiological apoptotic pathways using Bcl-2 or Bcl-xL inhibitors, or promoting the cell competition pathway described here, may have strikingly beneficial effects.

## Acknowledgements

We thank to Troy Littleton, Sergio Casas-Tintó, Richard Baines, the Vienna Drosophila Resource Center, the Bloomington Stock Center and the Developmental Studies Hybridoma Bank for sending stocks and reagents. We also acknowledge Julien Colomb for providing equipment and software (CeTrAn V4) used in the walking behaviour assay. We are grateful to the fly community at Champalimaud Research for criticial feedeback and for sharing antibodies and fly stocks. We thank technicians of the Champalimaud Fly Platform for support with the fly work and stock maintenance and Gil Costa for helping with scientific drawings. D.S.C. was supported by an EMBO long-term fellowship (ALTF 979-2014). Work in our laboratory is funded by the European Research Council, the Swiss National Science Foundation, the Portuguese Science Foundation, the Josef Steiner Cancer Research Foundation and the Swiss Cancer League.

## Contributions

D.S.C., C.R. and E.M. designed the experiments. D.S.C. performed and analysed the experiments. C.R. obtained and analysed data shown in Fig.6. S.S. obtained and analysed the data shown in Fig.7. M.M.M obtained preliminary data. B.H., B.T. and C.T. generated the molecular biology reagents. D.S.C., C.R. and E.M. wrote the manuscript.

## STAR methods

### Contact for Reagent and Resource Sharing

Further information and requests for resources and reagents should be directed to the Lead Contact, Eduardo Moreno

### Key Resources Table

| Antibodies | Source | Identifier |
| --- | --- | --- |
| rat anti-ELAV | DSHB | 7E8A10 |
| polyclonal anti-Myc-tag | Cell Signaling | 2272 |
| monoclonal mouse anti-Aβ42 | Synaptic Systems | 218 721 |
| anti-Fasciclin II | DSHB | 1D4 |
| chicken anti-GFP | Abcam | ab13970 |
| phalloidin Alexa-546 | Molecular Probes | A22283 |
| cleaved DCP1 | Cell Signaling | 9578 |
| Chemicals | Source | Identifier |
| Terminal Transferase, recombinant (rTdT) | Roche | 3333574001 |
| Biotin-16-dUTP) | Roche a | 11 093 070 910 |
| RU486 (Mifepristone) | Sigma | M8046-1G |
| propidium iodide | Sigma | P4170 |
| proteostat aggresome dye | Enzo Life Sciences | ENZ-51035-0025 |
| Fly stocks | Source | Identifier |
| *UAS-Aβ42 (2x)* | Casas-Tinto et al., 2011 | NA |
| *GMR-Gal4* | Bloomington | 1104 |
| *elav-GeneSwitch* | Bloomington | 43642 |
| *azot{KO; GFP}* | Merino et al., 2015 | NA |
| *azot{KO; w+}* | Merino et al., 2015 | NA |
| *UAS-diap1 (third,)* | Bloomington | 6657 |
| *UAS-flower<sup>LoseA</sup>* | Rhiner et al., 2010 | NA |
| *flower<sup>LoseA/B</sup> RNAi (long hairpin)* | Merino et al., 2013 | NA |
| *UAS-flower RNAi kk* | VDRC | 104993 |
| *UAS-RNAi azot GD* | VDRC | 18166 |
| *UAS-RNAi azot kk* | VDRC | 102353 |
| *UAS-secEMAP* | Casas-Tintó et al., 2015; | NA |
| *UAS-Aβ40* | Bloomington | #64215 |
| *UAS-httQ0* and *UAS-httQ128* | Lee et al., 2004 | NA |
| *UAS-αsynWT* and *UAS-αsynA30P* | Bloomington | #8146<br>#8147 |
| *flower<sup>ubi</sup>-YFP, flower<sup>LoseA</sup>-GFP, flower<sup>LoseB</sup>-RFP, and flower<sup>ubi</sup>-flag, flower<sup>LoseA</sup>-HA, flower<sup>LoseB</sup>-myc* | Yao et al., 2009 | NA |
| *UAS-lacZ* | Bloomington | 3955 |

## Experimental Model and Subject Details

### Fly husbandry and stocks

Flies were maintained on standard cornmeal-molasses-agar media and reared at 25°C under 12h alternating light-dark cycles. Stocks used in this study are listed in the Key Resources Table.

## Method Details

### Generation of *flower^LoseB^::mCherry* and *azot::mCherry reporter*

The *flower^LoseB^::mCherry knock-in* was made by genomic engineering (Baena-Lopez et al., 2013; Huang et al., 2009). The genomic engineering by Huang is a 2-step process consisting of ends-out gene targeting followed by phage integrase *phi31*-mediated DNA integration. A founder knock-out line was established with a genomic deletion of the *flower* locus at position 3L: 1’5816’737-15810028. A *knock-in* construct containing the deleted *flower* locus fused with mCherry after exon 6 (specific for *flower^LoseB^* isoform) was integrated in the KO line. The knock- in construct was done by site directed mutagenesis to remove the stop codon of exon 6 and add a restriction site to clone mCherry. The *knock-out* of *flower* and the *knock-in flower^LoseB^::mCherry* were proven by PCR. Vectors used for generating the *flower^LoseB^::mCherry* were as following: *pGX-attP*: (Knock-out vector), *pGEM-T* (used for the site directed mutagenesis and insertion of mCherry) and *pGEattB^GMR^* (*Knock- in* vector).

Primers used were the following: AAGCGGCCGCAGCAGCAACAACAGCAGCAACG and AAGCGGCCGCACCGTTCAATATGCAGGCGGC (5’ arm *flower* amplification), GGAGATCTGGATGATTCCTGAGCTGCGGTAT and AACTGCAGATGGGGACACCTAAAGAGGCACC (3’ arm *flower* amplification),

AACTATATTGGGCCGGCCAAGCTAACCGAATGCAAGAGGAACCGGAACCTA and GCATTCGGTTAGCTTGGCCGGCCCAATATAGTTTCTCACTAAAAATATATGCTTGC (mutagenesis primers), TAGGGCCGGCCATGGTGTCCAAGGGCGAAG and ATGGCCGGCCCTTATTTATACAGCTCGTCCATGC (mCherry amplification), ACATAGATCTATAAAAGCTTTCAATGTACACAAATTTG and AGCTGGCGCGCCAAAAAGCATGCCCCACAATAGTTAC (for *flower* KI).

The *azot::mCherry* reporter was generated by fusion PCR to combine *mCherry* coding sequence to *azot* genomic region, including 2430bps upstream of the start codon, the full *azot* exon and 175 bp at the 3’ UTR. *azot* genomic region was amplified from the Bac clone CH321-21G13 (http://pacmanfly.org/) using the following primers: TTGCTTAGACTGTGGCCAGAG and CTCTTCGCCCTTGGACACCATTCGCATTGTCATCATGTTGACGA for 5’ region and *azot* exon; GACATCTTCTCGCCCAGGTTG and ATGGACGAGCTGTATAAATAACCTCCATGTGAGTACTCGTA for 3’ UTR; GAGATCTCGACGTTCATACGGACGGACAGGCAGACGGAAGGAC and ACTGCATATAACATGCGCGAGA for the promoter region of *azot*. *mCherry* was amplified from *c5_stable2_neo* vector with primers TCAACATGATGACAATGCGAATGGTGTCCAAGGGCGAAGAG and ACGAGTACTCACATGGAGGTTATTTATACAGCTCGTCCATG. The final construct was obtained by two rounds of fusion PCRs (first with primers TAGGCGCGCCCCGCTCATTGTTTCCAAAGTGATTTTC and GCCGCTAGCGTATGAACGTCGAGATCTCGG; second with primers ACTGCATATAACATGCGCGAGA and TAGGCGCGCCCCGCTCATTGTTTCCAAAGTGATTTTC) and was cloned in *pGEattBGMR* with the restriction sites *NheI* and *AscI*.

### Immunohistochemistry and image acquisition

Wandering third instar larvae were collected and eye imaginal discs dissected. For clone induction, larvae were given a heat shock at 37ºC 48h or 72h before dissection. For pupal dissections, white prepupae (0hr) were collected and maintained at 25°C for 40h. Dissections were performed in chilled PBS, samples were fixed for 30min in *formaldehyde* (4% v/v in *PBS)* and permeabilized with PBT 0,4%Triton.

The primary antibodies and fluorescent reagents used in this study are listed in the key resource table. TUNEL staining (Roche) was performed according to the supplier’s protocol and modified as previously (Lolo et al., 2012). For detection of protein inclusions, brains were fixed and permeabilized as described above and incubated for 1h30min with the proteostat aggresome dye (Enzo Life Sciences) before mounting. For the necrosis assay, brains were dissected in PBS 1X and incubated for 30min at room temperature with 10 μg/ml propidium iodide (PI) (Sigma-Aldrich) in Schneider medium, following by washing and standard fixation ((Liu et al., 2014; Yang et al.,2013)). Samples were mounted in Vectashield (Vectorlab) and imaged on a Leica confocal SP5 or a Zeiss LSM 880 using a 20X dry objective or a 40X oil objective.

### Longevity Assays, brain morphology and neuronal manipulation

To minimize disturbing neural development and reduce differences on the genetic background, the RU486-inducible *GeneSwitch* system was employed (Osterwalder et al., 2001; Roman et al., 2001). The stock solution of RU486 (Mifepristone, Sigma, prepared in 80% ethanol) was diluted in MiliQ water to a final concentration of 100μM and 300μl of the diluted solution was added to the surface of the fly food and allowed to dry at room temperature for 48-36h (Poirier et al., 2008). For the mock solution, 80% ethanol was diluted 10X in water. For survivorship analysis, newly eclosed flies were transferred to bottles and allowed to mate for 2 days at 25ºC (He and Jasper, 2014; Linford et al., 2013). Females were then sorted into groups of 15-20 (more than 100 flies in total were used per genotype) and placed at 29ºC into vials containing standard food supplemented either with RU468 or mock solution. Flies were transferred to new food every 2-3 days and dead/censored animals were counted. For brain morphology analysis, males were subjected to the same protocol, aged at 29ºC until the required stage and dissected. To quantify number of *azot*-expressing cells with *azot{KO;GFP},* newly eclosed males were collected and kept at 25ºC to age for 5, 15 or 30 days.

### Behavioral assays

Detailed protocol and further description of the Buridan’s arena can be found elsewhere (Colomb and Brembs, 2014; Colomb et al., 2012). Shortly, 2 weeks-old females, kept in a 12/12 hours light/dark regime, were raised on standard or RU486-containing food. The day before measurements, flies were CO_2_ -anaesthetized (max 5 min) and their wings were cut with surgical scissors at two thirds of their length. For recordings, flies were placed in the center of the Buridan’s platform in a dark room. The walking activity of each individual fly was recorded for 5 minutes with the Buritrack software (http://buridan.sourceforge.net). Individual tests were re-initalized when flies jumped from the platform or exhibited grooming behaviour. Walking behaviour was analyzed with the CeTrAn software V4 (https://github.com/jcolomb/CeTrAn/releases/tag/v.4).

Long-term memory (**Courtship suppression assay):** The repeat training courtship assay was used to assess 24 hour long-term memory formation as published previously (Fitzsimons et al., 2013). Briefly, a training session was conducted by coupling individual males with a freshly mated female for a period of seven hours, while sham males were housed alone and served as controls to verify that courtship activity of a specific genotype was intact. Males were induced on RU486 food for one week, which was previously shown to be sufficient to induce the *elavGS* driver and elicit *Aβ42* expression in fly heads (Rogers et al., 2012). After 24 hours, all males, trained and sham, were coupled with new mated females and courtship activity was measured over a period of ten minutes as the percentage of time spent courting (courtship index, CI)(Reza et al., 2013). A memory index (MI) was then calculated as the ratio between the CI of every trained male and the mean value of the CI of the sham males of the same genotype. A range of scores between zero and one was obtained, with zero indicating good memory and one indicating memory similar to a sham control(Ejima and Griffith, 2007), e.g. no memory. Normal memory is generally characterized by a MI of 0.5-0.7 (Fitzsimons et al., 2013). Collected flies were flipped onto fresh food every two days and kept at 25°C in a 12 hr light/dark cycle. *elavGS>Aβ* flies on standard food served as uninduced control. In all experiments, the experimenter was blind to the genotype of the flies. Experiments were performed under ambient light at 25°C with 65-70% relative humidity and recorded for 10min using a camcorder (Sony Handycam HDR PJ410).

### Image analysis and Statistics

Image quantification was done with Fiji. The number of positive cells in the adult brain for DCP1, TUNEL, Flower^LoseB^::mCherry, Flower^LoseB^::RFP or Azot::mCherry was counted on 40-µm-wide maximum projections including the anterior part of the optic lobe. Noise signal was removed using a Gaussian blur filter (sigma =1) and/or applying a background subtraction (rolling=20). GFP expressing cells in *azot{KO;GFP*} flies were assumed to be GFP-positive particles wider than 9pixels on a 25μm-thick projection (showing a 141μm^2^ field) of the optic lobe. Measure of death induction in eye imaginal discs was done by counting the number of TUNEL positive particles in 10μm-thick maximum projections. Spaces between phalloidin staining with an area >25μm2 were assumed to be neurodegenerative vacuoles. Presence of vacuoles was quantified two weeks after eclosion at a 10μm deep ventral plane located in the central brain (next to the mushroom body).

**Survival curves:** For statistical analysis, a log-rank test (Mantel Cox) was applied to determine significant differences between survival curves.

**Walking behaviour data** was analyzed by an ANOVA model, which was validated posthoc with Tukey-Anscombe plot and QQ plot of the residuals. p values were calculated comparing all experimental genotypes with each other and corrected for multiple testing using Holm’s method (Holm, S, 1979). The variables distance, activity time, and turning angle were chosen for analysis based on previous test experiments.

**Courtship suppression assay**, raw data was subjected to arcsine transformation in order to obtain a normal distribution and the memory indexes of each genotype were subjected to a one-way ANOVA followed by Bonferroni and Holm’s correction by comparing genotypes to *elavGS>Aβ* induced controls. When comparing only two genotypes, the Student’s T-test (two-tailed, unpaired) was used. Significance was set at P < 0.05.

The distribution of the number of positive cells (for DCP1, Flower^LoseB^::mCherry, Flower^LoseB^::RFP or Azot::mCherry) in the optic lobes of adult flies was analyzed for statistical significant differences between groups with a Kruskal-Wallis test and a Dunn’s test was applied for multiple comparisons between genotypes. The number of brain vacuoles per hemisphere was analyzed for homogeneity between genotypes with a Levene test and p-values were calculated with one way ANOVA and a Dunnett’s posthoc test. To determine statistical differences between genotypes for the number of TUNEL positive cells in eye discs, a one-way ANOVA test, followed by a Dunnett’s posthoc were applied. When only two groups were compared and data did not follow a normal distribution assessed by d’Agostino-Pearson omnibus test, statistical significance was accessed with a Mann-Whitney U non-parametric test (for example in the quantification of TUNEL positive cells in the adult brain, GFP positive cells in the optic lobe and clone area). The number of PH3 positive cells was analyzed with an unpaired t-test with Welch’s correction. All graphs are displayed as mean ± standard error.

## Supplemental Figure legends

**Figure S1.**
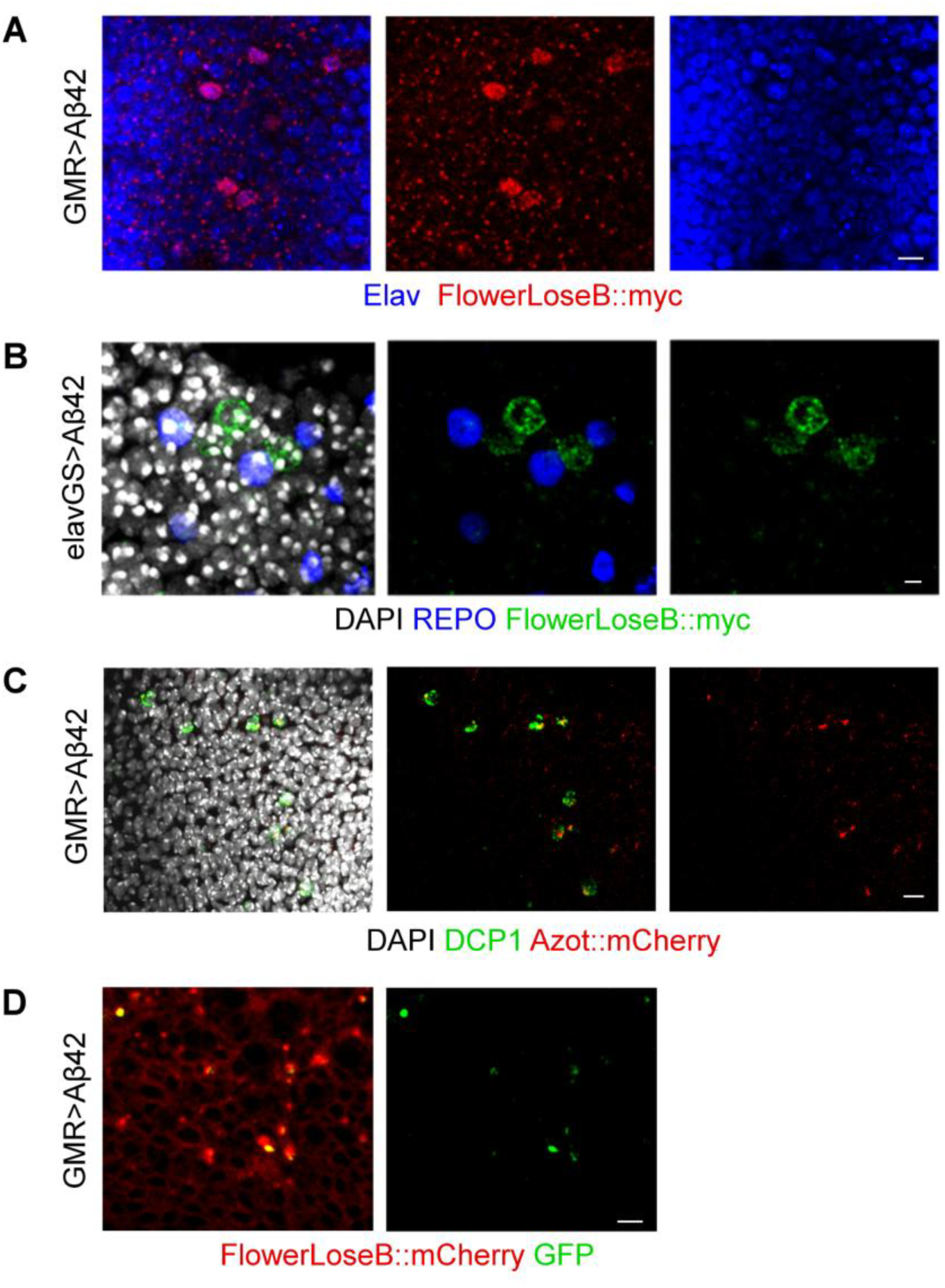
Neurons but not glial cells co-localize with Flower^LoseB^. Related to Figure 1 and Figure 4. **(A)** The Myc-tag (red) of the *flower^ubi^-flag, flower^LoseA^-HA, flower^LoseB^-myc* reporter co-localizes with neuronal cells marked by Elav (blue) in the optic lobe of *GMR>Aβ42* individuals. Scale bar: 5µm. **(B)** The Myc tag (green) of the flower^ubi^-flag, flower^LoseA^-HA, flower^LoseB^-myc reporter does not co-localize with REPO positive cells corresponding to glia (blue) in the optic lobe of *GMR>Aβ42* individuals. DAPI is in white. Scale bar: 2µm. **(C)** A subset of DCP1 labeled apoptotic cells (green) are also Azot::mCherry positive (red) in eye imaginal discs of *GMR>Aβ42* third instar larva. DAPI is in white. Scale bar: 5µm. **(D)** All GFP signal derived of the *azot{KO;GFP}* construct (green) co-localizes with Flower^LoseB^::mCherry cells (red), but not all Flower^LoseB^::mCherry expression co-localizes with GFP of azot{KO;GFP} in eye imaginal discs of *GMR>Aβ42* third instar larva. Scale bar: 5µm.

**Figure S2.**
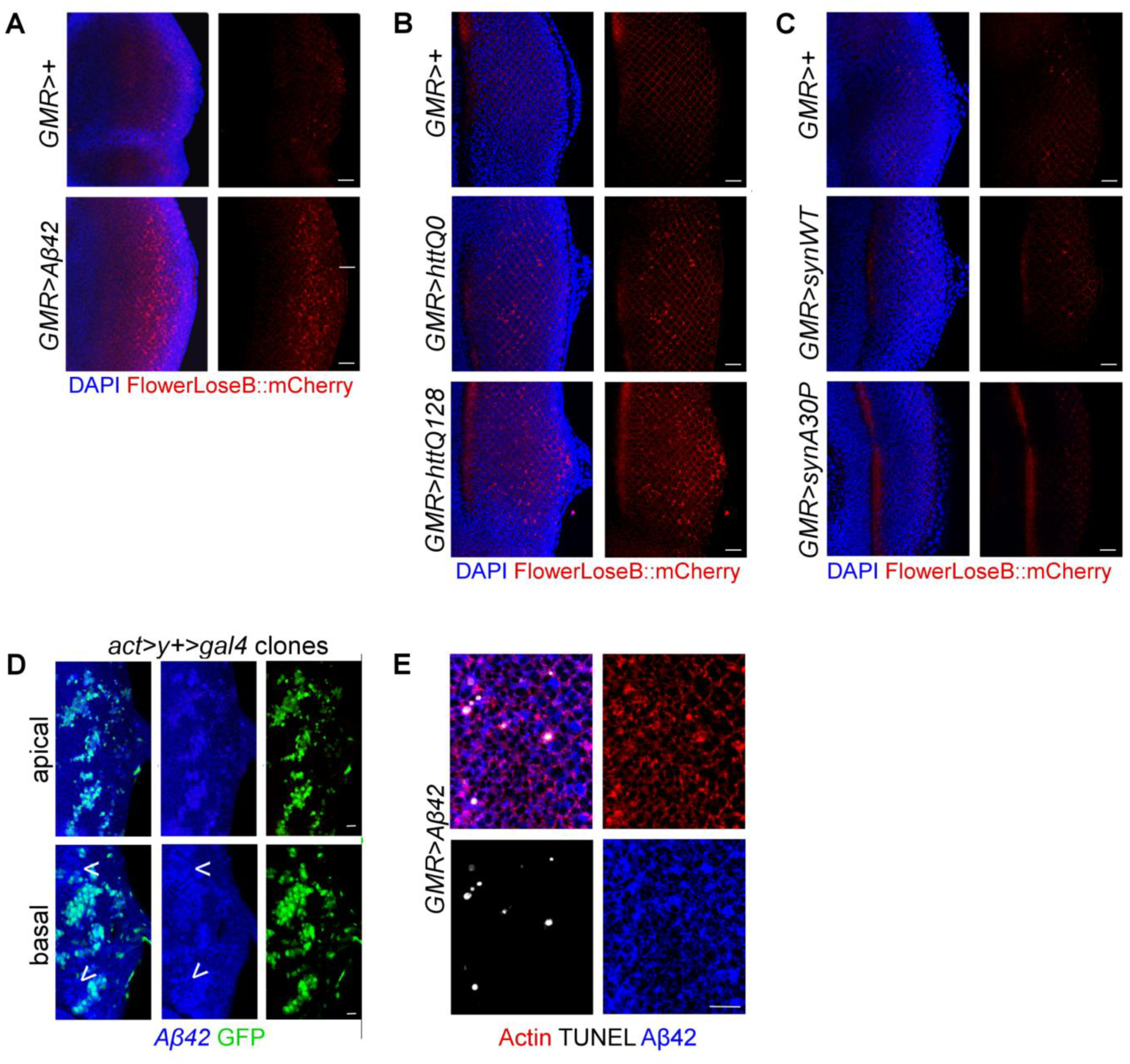
Flower^loseB^::mCherry reporter (red) is strongly induced by overexpression of *httQ128* in eye imaginal discs. Related to Figure 2 and Figure 3 **(A)** The Flower^loseB^::mCherry reporter (red) is strongly upregulated in *GMR>*Aβ42 eye imaginal discs of third instar larva comparing to *GMR>+* only control eye discs; the nuclear marker DAPI is shown in blue. Scale bar: 20 µm **(B)** The Flower^loseB^::mCherry reporter (red) is strongly expressed in *GMR>httQ128* eye imaginal discs, moderately expressed in *GMR>httQ0* eye discs but is not detected in *GMR>+* only controls; the nuclear marker DAPI is shown in blue. Scale bar: 20 µm **(C)** The Flower^loseB^::mCherry reporter (red) is not expressed in eye imaginal discs of *GMR>α-synWT* and *GMR> α-synA30P* larva. DAPI is shown in blue. Scale bar: 20 µm **(D)** Aβ42 localization (blue) in the apical side and basal side of eye discs 24h after the induction of *act>y+>gal4* clones (green). Arrows show Aβ42 accumulation out of the clone borders, evident on the basal side. Scale bar:10µm. **(E)** TUNEL labeling of cell death (white) detected in the eye disc of *GMR>Aβ42* larva. Actin is shown in red and Aβ42 peptide recognized by a specific antibody in blue. Scale bar: 10µm

**Figure S3.**
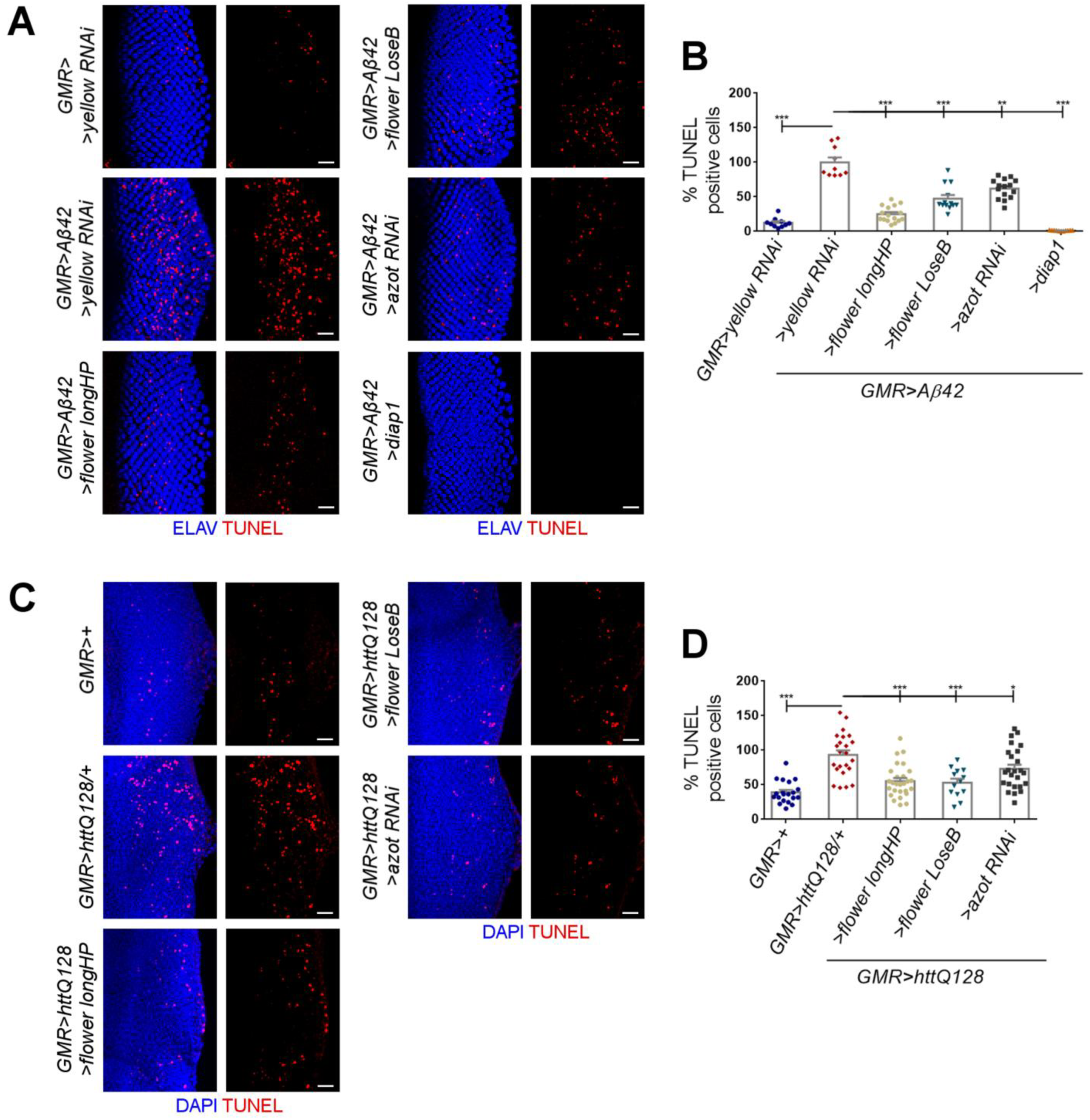
*flower* and *azot* mediate cell death induced by toxic proteins related to neurodegenerative diseases. Related to Figure 4 **(A)** Eye imaginal discs of third instar larva showing TUNEL labeling (red) of apoptotic cells, representative for each genotype. The nuclear marker ELAV is in blue. Scale bar: 20µm cells **(B)** Quantification of the number of TUNEL-positive cells per eye disc, assuming the number of apoptotic cells in *GMR>Aβ42*/ *yellow RNAi* as 100%. **(C)** Eye imaginal discs of third instar larva showing TUNEL labeling (red) of apoptotic cells, representative for each genotype. The nuclear marker DAPI is in blue. Scale bar: 20 µm **(D)** Quantification of the number of TUNEL-positive cells per eye disc, assuming the number of apoptotic cells in *GMR>httQ128* / *+* as 100%. Error bars show SEM. ***P value<0,001. **P value<0,01. *P value<0,05.

**Figure S4.**
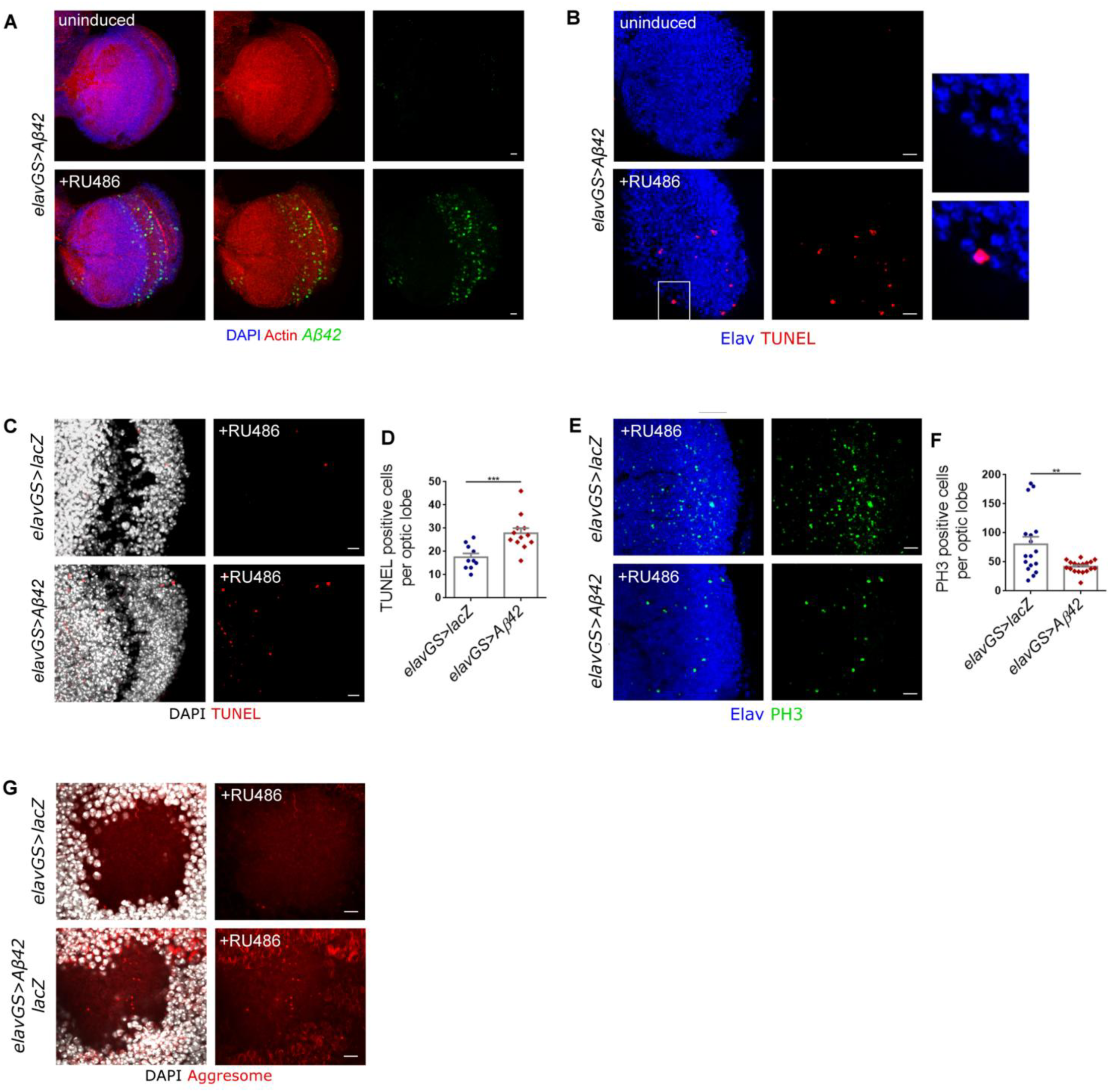
Conditional expression of human Aβ42 in the adult brain of *Drosophila* causes neuronal death and the formation of inclusion bodies. Related to Figure 5. **(A)** Conditional expression of human Aβ42 peptide in the *Drosophila* adult brain under the control of the *elav-GeneSwitch (elavGS)* driver in the presence of RU486. Aβ42 aggregates (green) are detected in the optic lobe of induced *elavGS>Aβ42* flies but not in uninduced controls. Actin is in red and nuclei are in blue. Scale bar:10 µm. **(B)** TUNEL-positive cells (red) in the optic lobes of *elavGS>Aβ42* males, uninduced or fed with RU486 for 10days. Neurons are marked by ELAV in blue. Scale bar: **(C,D)** Quantification and representative images of apoptotic cells in the optic lobe of *elavGS>lacZ* or *elavGS>Aβ42* flies fed on RU486-supplemented food. Apoptotic cells are labelled by TUNEL (red) and DAPI is in white. Scale bar: 5µm. **(E)** PH3 staining (green) in the optic lobe of e*lavGS>lacZ* or elavGS>Aβ42 flies treated with RU486. Elav is in blue. Scale bar: 10µm. **(F)** Quantification of the number of PH3 positive cells per optic lobe of the indicated genotypes at two weeks old. 10µm **(G)** Inclusion bodies formed by aggregated proteins (or aggresomes - in red) detected by the proteostat aggresome dye located in the mushroom body calyx of *elavGS>lacZ* or e*lavGS>Aβ42 / >lacZ* flies raised on food supplemented with RU486. DAPI in white. Scale bar: 5µm Error bars show standard error mean. ***P value<0,001, **P value<0,01. *P value<0,05.

**Figure S5.**
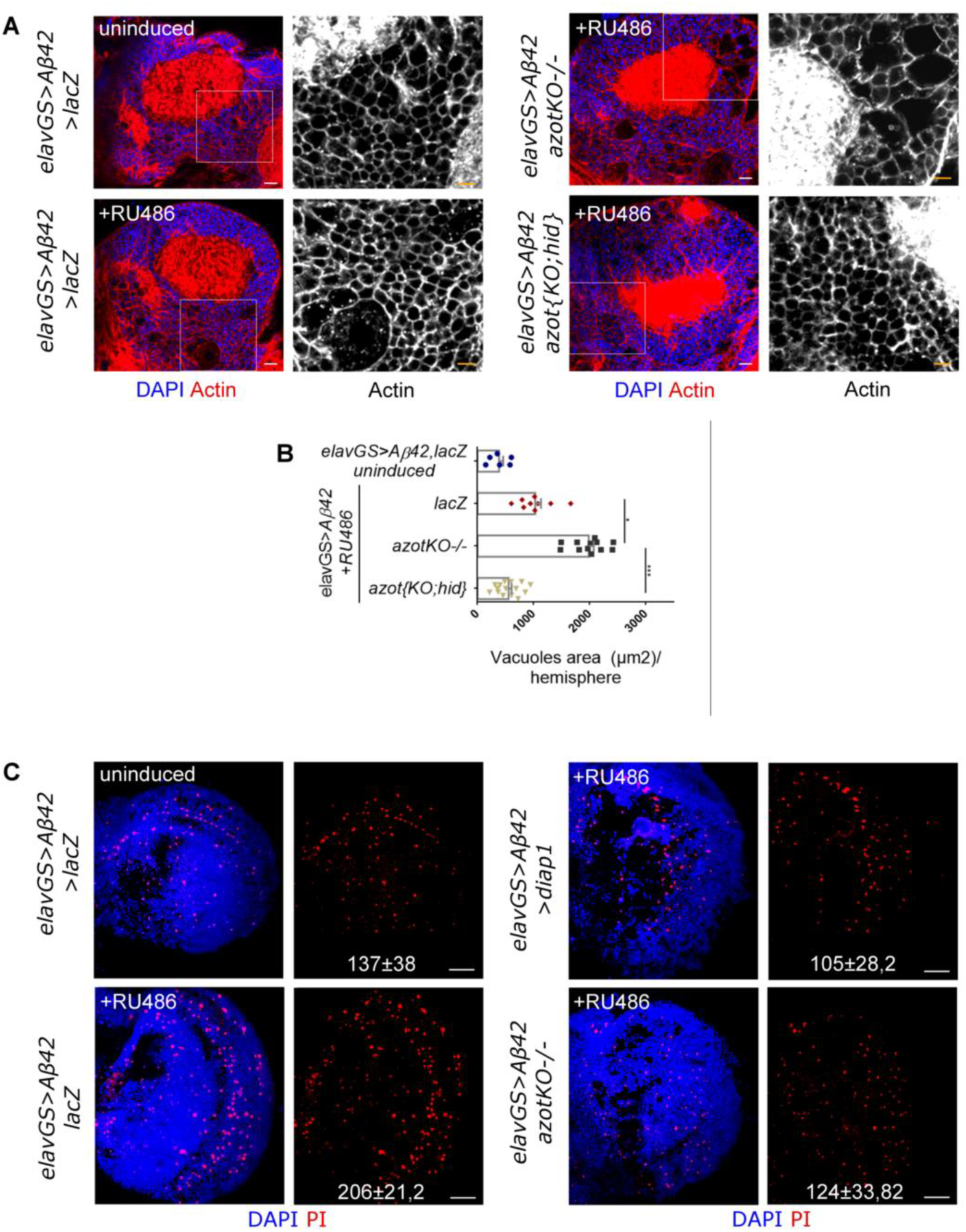
Azot is exclusively dedicated to cell death during neurodegeneration. Related to Figure 5. **(A)** Ventral plane of the central brain, focusing on the region adjacent to the mushroom body to show degenerative vacuoles present in the following genotypes: uninduced *elavGS>Aβ42/ >lacZ*, induced *elavGS>Aβ42/>lacZ* (+RU486), induced *elavGS>Aβ42, azotKO-/-* (+RU486), induced *elavGS>Aβ42/azot{KO;hid}* (+RU486). Phalloidin binding actin filaments is in red and DAPI shows nuclei in blue. Scale bar: 20µm in color pictures or 10µm in grayscale insets. **(B)** Mean of the total area occupied by vacuoles (in µm^2^) per brain section (at a 10µm deep plane). Error bars show SEM. ***P value<0,001, *P value<0,05. **(C)** Image displaying necrotic cells marked by PI (red) in the optic lobe of two weeks old males, representative for each genotype: uninduced *elavGS>Aβ42/ >lacZ*, induced *elavGS>Aβ42/ >lacZ* (+RU486), induced *elavGS>Aβ42, azotKO^-/-^* (+RU486) and induced *elavGS>Aβ42/ >diap1.* The average number (and standard deviation of the mean) of positive PI cells per optic lobe for each genotype is shown at the bottom of the image. DAPI is in blue. Scale bar: 20µm.

**Figure S6.**
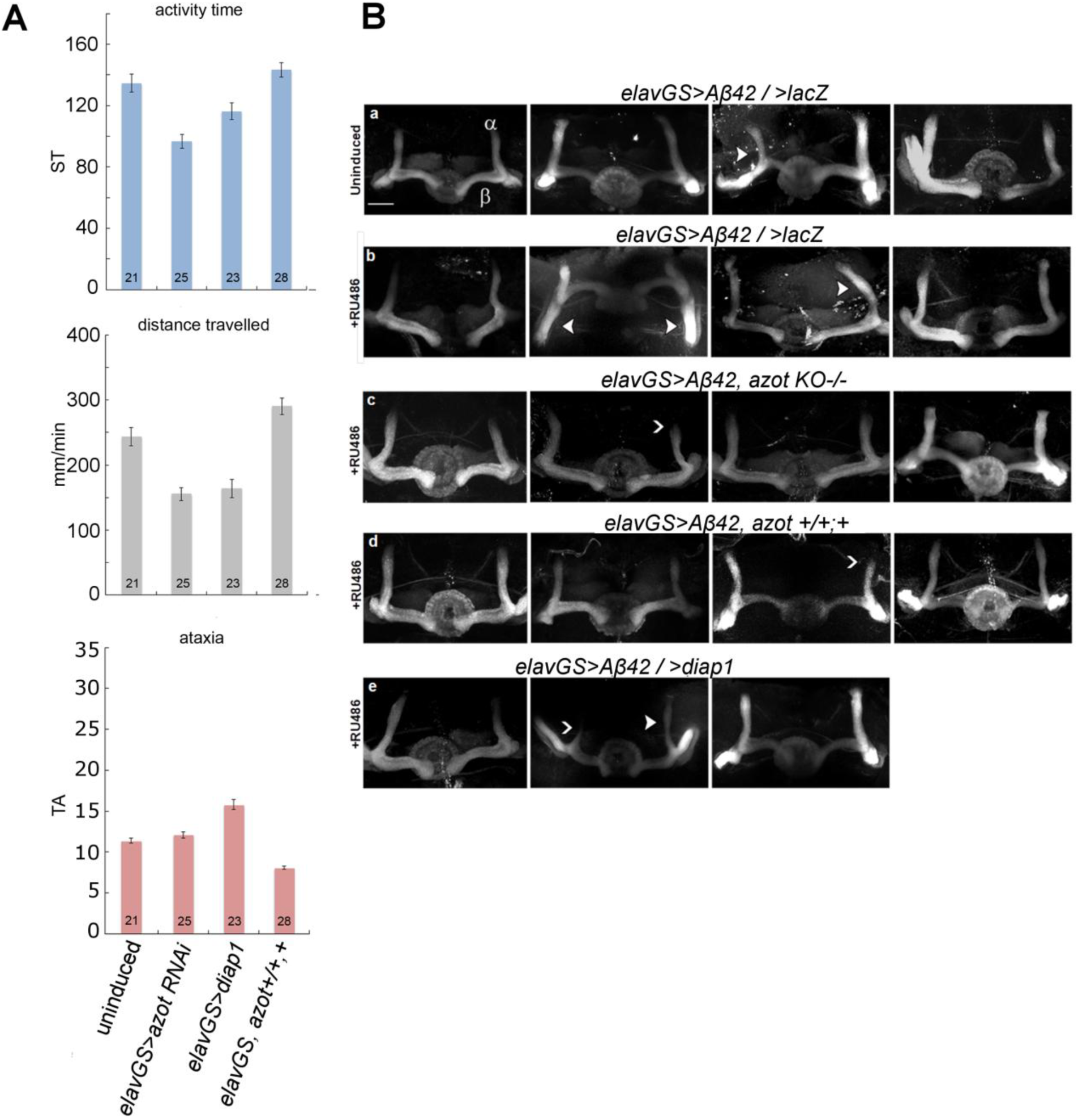
Morphology of the mushroom body (α and β lobes). Related to Figure 6. **(A)** Graphs depicting activity time in seconds (using a speed threshold, ST)(Colomb et al., 2012), distance walked in mm/min and median turning angle (TA) in degrees. Parameters were calculated from individual walks of two weeks-old heterozygous flies raised on RU486 for each genotype. Error bars show S.E.M., numbers indicate number of individual flies tested. **(B)** Immunohistochemistry with anti-FasII on whole-mount brains depicting the morphology of the mushroom body lobes in one week-old males of the following genotypes: uninduced *elavGS>Aβ42/ >lacZ*, induced *elavGS>Aβ42/>lacZ*, induced *elavGS>Aβ42, azotKO-/-*, induced *elavGS>Aβ42, azot+;+/+* and induced *elavGS>Aβ42/ >diap1*). A mild phenotypic variability of the mushroom body lobes was observed consisting of the following traits: outgrowth and guidance defects indicated by a white arrow and irregular shape indicated by a white arrowhead. Images are projections of confocal z-stacks. Scale bar = 50 μm.

## Supplemental Videos legends

**Buridan arena**

**Video 1 Locomotion of an uninduced *elavGS>Aβ42 / >lacZ* individual.**

Walking trajectories of heterozygous e*lavGS>Aβ42/+* females after two weeks on normal food.

**Video 2 Locomotion of an induced *elavGS>Aβ42 / >lacZ* individual.**

Walking behavior of heterozygous e*lavGS>Aβ42/+* females after two weeks on RU486.

**Video 3 Locomotion of *elavGS>Aβ42, azot+/+;+* individual.**

Walking trajectories of heterozygous *elavGS>Aβ42, azot+/+;+* females after two weeks on RU486.

**Courtship suppression assay**

Representative 3 min extracts of 10 min long videos recorded to evaluate long-term memory 24 hours after exposure to unreceptive female (trained) or no female in the same test chamber (sham controls).

**Videos 4** Courtship activity of trained e*lavGS>Aβ42 / >lacZ* flies, induced for one week on RU486.

**Videos 5**

Courtship activity of trained e*lavGS>Aβ42 / >lacZ* flies raised for 1 week on normal food (no induction).

**Videos 6**

Courtship activity of trained e*lavGS>Aβ42, azotKO^-^/^-^* flies, induced for one week on RU486.

**Video 7**

Courtship activity of trained e*lavGS>Aβ42 / >diap* fly induced for one week on RU486.

**Video 8**

Courtship activity of trained elavGS>Aβ42, azot+/+;+ fly induced for one week on RU486.

## References

Alexander, D.B., Ichikawa, H., Bechberger, J.F., Valiunas, V., Ohki, M., Naus, C.C.G., Kunimoto, T., Tsuda, H., Miller, W.T., and Goldberg, G.S. (2004). Normal cells control the growth of neighboring transformed cells independent of gap junctional communication and SRC activity. Cancer Res. 64, 1347–1358.

Ashe, K.H., and Zahs, K.R. (2010). Probing the biology of Alzheimer’s disease in mice. Neuron 66, 631–645.

Baena-Lopez, L.A., Alexandre, C., Mitchell, A., Pasakarnis, L., and Vincent, J.-P. (2013). Accelerated homologous recombination and subsequent genome modification in Drosophila. Development 140, 4818–4825.

Braak, H., and Braak, E. (1991). Neuropathological stageing of Alzheimer-related changes. Acta Neuropathol. (Berl.) 82, 239–259.

Casas-Tinto, S., Zhang, Y., Sanchez-Garcia, J., Gomez-Velazquez, M., Rincon-Limas, D.E., and Fernandez-Funez, P. (2011). The ER stress factor XBP1s prevents amyloid-neurotoxicity. Hum. Mol. Genet. 20, 2144–2160.

Casas-Tintó, S., Lolo, F.-N., and Moreno, E. (2015). Active JNK-dependent secretion of Drosophila Tyrosyl-tRNA synthetase by loser cells recruits haemocytes during cell competition. Nat. Commun. 6, 10022.

Colomb, J., and Brembs, B. (2014). Sub-strains of Drosophila Canton-S differ markedly in their locomotor behavior. F1000Research.

Colomb, J., Reiter, L., Blaszkiewicz, J., Wessnitzer, J., and Brembs, B. (2012). Open Source Tracking and Analysis of Adult Drosophila Locomotion in Buridan’s Paradigm with and without Visual Targets. PLoS ONE 7, e42247.

de la Cova, C., Abril, M., Bellosta, P., Gallant, P., and Johnston, L.A. (2004). Drosophila myc regulates organ size by inducing cell competition. Cell 117, 107–116.

Crittenden, J.R., Skoulakis, E.M., Han, K.A., Kalderon, D., and Davis, R.L. (1998). Tripartite mushroom body architecture revealed by antigenic markers. Learn. Mem. Cold Spring Harb. N 5, 38–51.

Eichenlaub, T., Cohen, S.M., and Herranz, H. (2016). Cell Competition Drives the Formation of Metastatic Tumors in a Drosophila Model of Epithelial Tumor Formation. Curr. Biol. CB 26, 419–427.

Ejima, A., and Griffith, L.C. (2007). Measurement of Courtship Behavior in Drosophila melanogaster. Cold Spring Harb. Protoc. 2007, pdb.prot4847.

Feany, M.B., and Bender, W.W. (2000). A Drosophila model of Parkinson’s disease. Nature 404, 394–398.

Fernández-Hernández, I., Rhiner, C., and Moreno, E. (2013). Adult neurogenesis in Drosophila. Cell Rep. 3, 1857–1865.

Fitzsimons, H.L., Schwartz, S., Given, F.M., and Scott, M.J. (2013). The Histone Deacetylase HDAC4 Regulates Long-Term Memory in Drosophila. PLoS ONE 8, e83903.

Fushima, K., and Tsujimura, H. (2007). Precise control of fasciclin II expression is required for adult mushroom body development in Drosophila. Dev. Growth Differ. 49, 215–227.

Gibson, M.C., and Perrimon, N. (2005). Extrusion and death of DPP/BMP-compromised epithelial cells in the developing Drosophila wing. Science 307, 1785–1789.

Gómez-Isla, T., Price, J.L., McKeel, D.W., Morris, J.C., Growdon, J.H., and Hyman, B.T. (1996). Profound loss of layer II entorhinal cortex neurons occurs in very mild Alzheimer’s disease. J. Neurosci. Off. J. Soc. Neurosci. 16, 4491–4500.

Hay, B.A., Wassarman, D.A., and Rubin, G.M. (1995). Drosophila homologs of baculovirus inhibitor of apoptosis proteins function to block cell death. Cell 83, 1253–1262.

He, Y., and Jasper, H. (2014). Studying aging in Drosophila. Methods 68, 129–133.

Hogan, C., Dupré-Crochet, S., Norman, M., Kajita, M., Zimmermann, C., Pelling, A.E., Piddini, E., Baena-López, L.A., Vincent, J.-P., Itoh, Y., et al. (2009). Characterization of the interface between normal and transformed epithelial cells. Nat. Cell Biol. 11, 460–467.

Huang, Y., and Mucke, L. (2012). Alzheimer mechanisms and therapeutic strategies. Cell 148, 1204–1222.

Huang, J., Zhou, W., Dong, W., Watson, A.M., and Hong, Y. (2009). Directed, efficient, and versatile modifications of the Drosophila genome by genomic engineering. Proc. Natl. Acad. Sci. 106, 8284–8289.

Igaki, T. (2009). Correcting developmental errors by apoptosis: lessons from Drosophila JNK signaling. Apoptosis Int. J. Program. Cell Death 14, 1021–1028.

Iijima, K., Liu, H.-P., Chiang, A.-S., Hearn, S.A., Konsolaki, M., and Zhong, Y. (2004). Dissecting the pathological effects of human A 40 and A 42 in Drosophila: A potential model for Alzheimer’s disease. Proc. Natl. Acad. Sci. 101, 6623–6628.

Kajita, M., and Fujita, Y. (2015). EDAC: Epithelial defence against cancer-cell competition between normal and transformed epithelial cells in mammals. J. Biochem. (Tokyo) 158, 15–23.

Karran, E., and De Strooper, B. (2016). The amyloid cascade hypothesis: are we poised for success or failure? J. Neurochem. 139 Suppl 2, 237–252.

Karran, E., Mercken, M., and De Strooper, B. (2011). The amyloid cascade hypothesis for Alzheimer’s disease: an appraisal for the development of therapeutics. Nat. Rev. Drug Discov. 10, 698–712.

Keleman, K., Krüttner, S., Alenius, M., and Dickson, B.J. (2007). Function of the Drosophila CPEB protein Orb2 in long-term courtship memory. Nat. Neurosci. 10, 1587–1593.

Kolahgar, G., Suijkerbuijk, S.J.E., Kucinski, I., Poirier, E.Z., Mansour, S., Simons, B.D., and Piddini, E. (2015). Cell Competition Modifies Adult Stem Cell and Tissue Population Dynamics in a JAK-STAT-Dependent Manner. Dev. Cell 34, 297–309.

Lee, W.-C.M., Yoshihara, M., and Littleton, J.T. (2004). Cytoplasmic aggregates trap polyglutamine-containing proteins and block axonal transport in a Drosophila model of Huntington’s disease. Proc. Natl. Acad. Sci. 101, 3224–3229.

Levayer, R., Hauert, B., and Moreno, E. (2015). Cell mixing induced by myc is required for competitive tissue invasion and destruction. Nature 524, 476–480.

Linford, N.J., Bilgir, C., Ro, J., and Pletcher, S.D. (2013). Measurement of Lifespan in *Drosophila melanogaster* J. Vis. Exp.

Liu, K., Ding, L., Li, Y., Yang, H., Zhao, C., Lei, Y., Han, S., Tao, W., Miao, D., Steller, H., et al. (2014). Neuronal necrosis is regulated by a conserved chromatin-modifying cascade. Proc. Natl. Acad. Sci. 111, 13960–13965.

Lolo, F.-N., Casas-Tintó, S., and Moreno, E. (2012). Cell competition time line: winners kill losers, which are extruded and engulfed by hemocytes. Cell Rep. 2, 526–539.

Martins, V.C., Busch, K., Juraeva, D., Blum, C., Ludwig, C., Rasche, V., Lasitschka, F., Mastitsky, S.E., Brors, B., Hielscher, T., et al. (2014). Cell competition is a tumour suppressor mechanism in the thymus. Nature 509, 465–470.

Menéndez, J., Pérez-Garijo, A., Calleja, M., and Morata, G. (2010). A tumor-suppressing mechanism in Drosophila involving cell competition and the Hippo pathway. Proc. Natl. Acad. Sci. U. S. A. 107, 14651–14656.

Merino, M.M., Rhiner, C., Portela, M., and Moreno, E. (2013). “Fitness fingerprints” mediate physiological culling of unwanted neurons in Drosophila. Curr. Biol. CB 23, 1300–1309.

Merino, M.M., Rhiner, C., Lopez-Gay, J.M., Buechel, D., Hauert, B., and Moreno, E. (2015). Elimination of Unfit Cells Maintains Tissue Health and Prolongs Lifespan. Cell 160, 461–476.

Merino, M.M., Levayer, R., and Moreno, E. (2016). Survival of the Fittest: Essential Roles of Cell Competition in Development, Aging, and Cancer. Trends Cell Biol. 26, 776–788.

Moreno, E., and Basler, K. (2004). dMyc transforms cells into super-competitors. Cell 117, 117–129.

Moreno, E., Basler, K., and Morata, G. (2002). Cells compete for decapentaplegic survival factor to prevent apoptosis in Drosophila wing development. Nature 416, 755–759.

Moreno, E., Fernandez-Marrero, Y., Meyer, P., and Rhiner, C. (2015). Brain regeneration in Drosophila involves comparison of neuronal fitness. Curr. Biol. CB 25, 955–963.

Nichols, C.D., Becnel, J., and Pandey, U.B. (2012). Methods to assay Drosophila behavior. J. Vis. Exp. JoVE.

Ossenkoppele, R., Cohn-Sheehy, B.I., La Joie, R., Vogel, J.W., Möller, C., Lehmann, M., van Berckel, B.N.M., Seeley, W.W., Pijnenburg, Y.A., Gorno-Tempini, M.L., et al. (2015). Atrophy patterns in early clinical stages across distinct phenotypes of Alzheimer’s disease. Hum. Brain Mapp. 36, 4421–4437.

Osterwalder, T., Yoon, K.S., White, B.H., and Keshishian, H. (2001). A conditional tissue-specific transgene expression system using inducible GAL4. Proc. Natl. Acad. Sci. 98, 12596–12601.

Petrova, E., López-Gay, J.M., Rhiner, C., and Moreno, E. (2012). Flower-deficient mice have reduced susceptibility to skin papilloma formation. Dis. Model. Mech. 5, 553–561.

Poirier, L., Shane, A., Zheng, J., and Seroude, L. (2008). Characterization of the Drosophila gene-switch system in aging studies: a cautionary tale. Aging Cell 7, 758–770.

Portela, M., Casas-Tinto, S., Rhiner, C., López-Gay, J.M., Domínguez, O., Soldini, D., and Moreno, E. (2010). Drosophila SPARC is a self-protective signal expressed by loser cells during cell competition. Dev. Cell 19, 562–573.

Reza, M.A., Mhatre, S.D., Morrison, J.C., Utreja, S., Saunders, A.J., Breen, D.E., and Marenda, D.R. (2013). Automated analysis of courtship suppression learning and memory in Drosophila melanogaster. Fly (Austin) 7, 105–111.

Rhiner, C., López-Gay, J.M., Soldini, D., Casas-Tinto, S., Martín, F.A., Lombardía, L., and Moreno, E. (2010). Flower Forms an Extracellular Code that Reveals the Fitness of a Cell to its Neighbors in Drosophila. Dev. Cell 18, 985–998.

Rogers, I., Kerr, F., Martinez, P., Hardy, J., Lovestone, S., and Partridge, L. (2012). Ageing Increases Vulnerability to Aβ42 Toxicity in Drosophila. PLoS ONE 7, e40569.

Roman, G., Endo, K., Zong, L., and Davis, R.L. (2001). P[Switch], a system for spatial and temporal control of gene expression in Drosophila melanogaster. Proc. Natl. Acad. Sci. U. S. A. 98, 12602–12607.

Saxena, S., and Caroni, P. (2011). Selective Neuronal Vulnerability in Neurodegenerative Diseases: from Stressor Thresholds to Degeneration. Neuron 71, 35–48.

Seab, J.P., Jagust, W.J., Wong, S.T., Roos, M.S., Reed, B.R., and Budinger, T.F. (1988). Quantitative NMR measurements of hippocampal atrophy in Alzheimer’s disease. Magn. Reson. Med. 8, 200–208.

Siegel, R.W., and Hall, J.C. (1979). Conditioned responses in courtship behavior of normal and mutant Drosophila. Proc. Natl. Acad. Sci. U. S. A. 76, 3430–3434.

Soldano, A., and Hassan, B.A. (2014). Beyond pathology: APP, brain development and Alzheimer’s disease. Curr. Opin. Neurobiol. 27, 61–67.

Song, L., He, Y., Ou, J., Zhao, Y., Li, R., Cheng, J., Lin, C.-H., and Ho, M.S. (2017). Auxilin Underlies Progressive Locomotor Deficits and Dopaminergic Neuron Loss in a Drosophila Model of Parkinson’s Disease. Cell Rep. 18, 1132–1143.

Speretta, E., Jahn, T.R., Tartaglia, G.G., Favrin, G., Barros, T.P., Imarisio, S., Lomas, D.A., Luheshi, L.M., Crowther, D.C., and Dobson, C.M. (2012). Expression in Drosophila of Tandem Amyloid β Peptides Provides Insights into Links between Aggregation and Neurotoxicity. J. Biol. Chem. 287, 20748–20754.

Suijkerbuijk, S.J.E., Kolahgar, G., Kucinski, I., and Piddini, E. (2016). Cell Competition Drives the Growth of Intestinal Adenomas in Drosophila. Curr. Biol. CB 26, 428–438.

Tamori, Y., and Deng, W.-M. (2013). Tissue repair through cell competition and compensatory cellular hypertrophy in postmitotic epithelia. Dev. Cell 25, 350–363.

Wakabayashi, K., Honer, W.G., and Masliah, E. (1994). Synapse alterations in the hippocampal-entorhinal formation in Alzheimer’s disease with and without Lewy body disease. Brain Res. 667, 24–32.

Yang, Y., Hou, L., Li, Y., Ni, J., and Liu, L. (2013). Neuronal necrosis and spreading death in a Drosophila genetic model. Cell Death Dis. 4, e723.

Yao, C.-K., Lin, Y.Q., Ly, C.V., Ohyama, T., Haueter, C.M., Moiseenkova-Bell, V.Y., Wensel, T.G., and Bellen, H.J. (2009). A synaptic vesicle-associated Ca2+ channel promotes endocytosis and couples exocytosis to endocytosis. Cell 138, 947–960.

Zanetti, O., Solerte, S.B., and Cantoni, F. (2009). Life expectancy in Alzheimer’s disease (AD). Arch. Gerontol. Geriatr. 49 Suppl 1, 237–243.

